# Unexpected Kif4a functions in adult regeneration encompass a dual role in neurons and in proliferative repair Schwann cells

**DOI:** 10.1101/2023.05.21.541636

**Authors:** Patrícia D. Correia, Bárbara M. de Sousa, Jesús Chato-Astrain, Joana P. Faria, Veronica Estrada, João B. Relvas, Hans W. Müller, Víctor Carriel, Frank Bosse, Sandra I. Vieira

## Abstract

Contrary to the adult central nervous system (CNS), the peripheral nervous system (PNS) has an intrinsic ability to regenerate that, among others, passes by expressing regeneration-associated genes such as kinesin family members. We here show that Kinesin family motor protein 4a (KIF4A), associated to neurodevelopmental disorders and thought for long to be only embryonically expressed, is highly abundant in axons and Schwann cells of adult rat CNS and rat and human PNS. Moreover, *Kif4a* is up-regulated in injured PNS neurons, being detected in their nuclei and regrowing axons, consistent with its functions as a chromokinesin and in the axonal transport of e.g. β1-integrin and L1CAM. Interestingly, *Kif4a* is also highly up-regulated in Schwann cells transdifferentiating into a proliferative repair phenotype at the injured distal nerve stumps. A role for *Kif4a* in cultured Schwann cells proliferation was confirmed, with *Kif4a* mRNA expression being ∼6-fold higher in proliferating versus growth-arrested Schwann cells, and *Kif4a* knockdown impairing Schwann cells’ proliferation. To our knowledge, this is the first description of KIF4A expression in adult nervous systems, up-regulation in neuroregeneration and pro-neuroregenerative roles, including promoting Schwann cells proliferation. KIF4A dual role in axonal regeneration, through neurons and glia, places as an attractive target for future neuroregeneration therapies.

## INTRODUCTION

Traumatic spinal cord injuries (SCI) are complex pathologies of the Central Nervous System (CNS), usually affecting motor, sensory and autonomic functions of the patients, and that result in low quality of life, lower life expectancy, and in expensive medical care^1^. In 2016, the global incidence of SCI was 13 per 100 000, and its prevalence was 27.04 million cases. In most countries, falls and traffic accidents are the leading causes of these injuries^1^. Although its management has improved, SCI are yet untreatable, mostly due to the very low mammalian CNS regenerative capacity, that further declines into adulthood. However, unlike the failed attempts of the injured CNS fibers to regenerate, that are hindered by a persistent inhibitory environment at the lesion area^2–8^, injured nerves of the peripheral nervous system (PNS) are able to regenerate significantly. Small peripheral nerve defects can heal spontaneously, whereas in severe peripheral lesions regeneration can be assisted by autografts or artificial scaffolds^9–12^.

Following traumatic CNS or PNS injuries, complex pathophysiological processes occur at the injury site, including metabolic, transcriptomic and morphologic alterations^13,14^. While the very early events of peripheral nerve injury (PNI) resemble those of SCI, differences quickly follow. Upon PNI, various cells undergo morpho-functional changes and switch from a maintenance state to a pro-regenerative one, in a process termed ‘Wallerian degeneration’^15^. At the proximal nerve stump, axons degenerate back for some distance from the injury site (‘retrograde degeneration’) leaving the endoneural tubes empty. This retraction causes Schwann cells distal to the lesion site to lose axonal contact, resulting in significant alterations in the signaling environment. Together with a cocktail of factors secreted by macrophages, these trigger Schwann cells transdifferentiation into a repair-promoting phenotype^16^. The ‘repair Schwann cells’ proliferate and activate pro-regenerative events, including myelin autophagy and up-regulation of macrophages-recruiting cytokines. These invading macrophages, alongside resident endoneurial macrophages, clean the remaining inhibitory myelin debris from the lesion site^17–21^. Repair Schwann cells further support the survival of injured neurons and axonal regeneration by expressing trophic factors and elongating and branching into regenerative tracks, necessary to guide axons back to their target areas (the ‘bands of Büngner’)^22^.

In contrast, the spinal cord does not restore functional connections after injury, due to a non-permissive environment. Oligodendrocytes, the counterpart of Schwann cells in the CNS, and other glial cells (reactive astrocytes and microglia) are less plastic and/or provide a post-injury milieu that minimizes the primary damage but prevents successful axonal regeneration.

Nevertheless, studies have shown that CNS neurons are to some extent capable of regeneration under certain circumstances^23–25^. For example, a peripheral nerve graft implanted into the SCI lesion site can trigger both axonal regeneration and the expression of regeneration-associated genes (RAGs) in central neurons ^23,25,26^. Considering that up-regulation of key RAGs in injured CNS neurons can enhance axonal regeneration^27^, better insight into the RAG program and network is crucial to identify regeneration promoters with therapeutic value for CNS lesions.

Genome-wide analyses aiming to identify RAGs are being conducted in neuroregenerative models, particularly adult PNI models^28^ and animals that naturally regenerate the spinal cord, such as *Xenopus laevis*^29^ and *Danio rerio*^30^. Injury-induced deregulation of several kinesin family members has been reported in such studies^28–32^ and, e.g., Kif3c has already been identified as an established RAG with important role for successful PNI/SCI recovery^33^. In one of our parallel studies, we have observed altered expression of *KIF4A* (kinesin family member 4a) levels in the supplementary regenerative datasets of these transcriptomic studies^29,34^. This was unexpected since KIF4A, a highly conserved ATP-dependent microtubule-based motor protein^35^ known to anterogradely transport cargo in developing neuronal axons^36^, has been reported to be absent or expressed at very low levels in the adulthood^37,38^. Current knowledge places KIF4A as a promoter of neuritogenesis and axon guidance during development^39^ by transporting key cargoes from the cytoplasm to axonal growth cones. These include the ribosomal protein P0 in developing dorsal root ganglia (DRG) neurons^40^, and the L1 neuritogenic neural cell adhesion molecules (CAM) of the immunoglobulin superfamily^40,41^ in hippocampal neurons^39^. Importantly, L1 protein (alias L1CAM) has already been related to regeneration of both adult central and peripheral neurons^26,42,43^. KIF4A also binds to and anterogradely transports other neuronal CAMs, like TAX1^39^ and β1-integrin, in immature neurons^38^. Additionally, like some other kinesin family members, KIF4A is also a chromokinesin with multiple nuclear roles. It supports chromosome condensation^44^ and binds to cell cycle-associated proteins to regulate mitotic spindle pre-cytokinesis organization, both important to maintain chromosome integrity and genomic stability^45^. KIF4A also plays a role in neuronal survival by suppressing PARP-1 enzymatic activity during neuronal development^46^, and its artificial overexpression in mature cultured neurons can cause massive apoptosis^38^, highlighting the need of tightly regulate KIF4A levels.

The importance of KIF4A role in developing nervous systems is emphasized by the association of abnormal KIF4A expression with various neuropathologies, such as developmental delay, intellectual disability, hydrocephalus and brain anomalies (manifestations of the ‘KIF4A-associated syndrome’)^47^, and other human cognitive phenotypes, like Mental Retardation X-linked 100, a developmental disorder characterized by microcephaly, cortical malformations, abnormal higher mental functions, seizures, and poor speech^47–49^ (www.malacards.org). *KIF4A* is also abnormally expressed in several cancers, including glioblastoma^50^, and its knockdown or overexpression can inhibit proliferation and migration of cancer cells.

Despite these known key neural developmental functions, nothing is known about the role of KIF4A in adult nervous systems. With this study we aimed to characterize this protein in adult central and peripheral neural tissues, and to further pursue a putative regeneration-associated role for KIF4A in adulthood. To this end, we used various models of sciatic nerve injury and regeneration, as well as functional assays in cultured rat Schwann cells. To our knowledge, this is the first description of KIF4A expression in adult nervous system, of its regeneration-associated regulation after injury, and of its role in Schwann cells proliferation.

## RESULTS

### KIF4A is expressed in adult rat spinal cord neurons, in basal state and after injury

To first analyze KIF4A distribution in the adult CNS, sagittal sections of the spinal cords of both sham-operated and injured animals were examined by immunohistochemistry (IHC). KIF4A-positive structures were present in both grey and white matter of the spinal cord (**Figure 1**). In the ventral grey matter, KIF4A was observed in glial fibrillary acidic protein (GFAP)-negative large cell bodies (including their nuclei), most likely motor neurons (**Figure 1A**, arrowheads). In the white matter, double labelling revealed overlap between the axonal neurofilament (NF) marker and KIF4A, confirming the neuronal distribution and highly abundant axonal expression of this protein (**Figure 1B**). Moreover, KIF4A appeared to be present in myelinated axons, as KIF4A-positive axons showed to be enclosed in structures positive for myelin basic protein (MBP), a CNS myelin marker (**Figure 1C**). Axonal staining of KIF4A occurs within myelin sheaths, supporting the idea that KIF4A is expressed in spinal cord axons. Co-labelling with the glial marker GFAP in white matter confirmed that there was no visible KIF4A/GFAP overlap in the intact spinal cord (**Figure 1A; D**).

**Figure 1.**
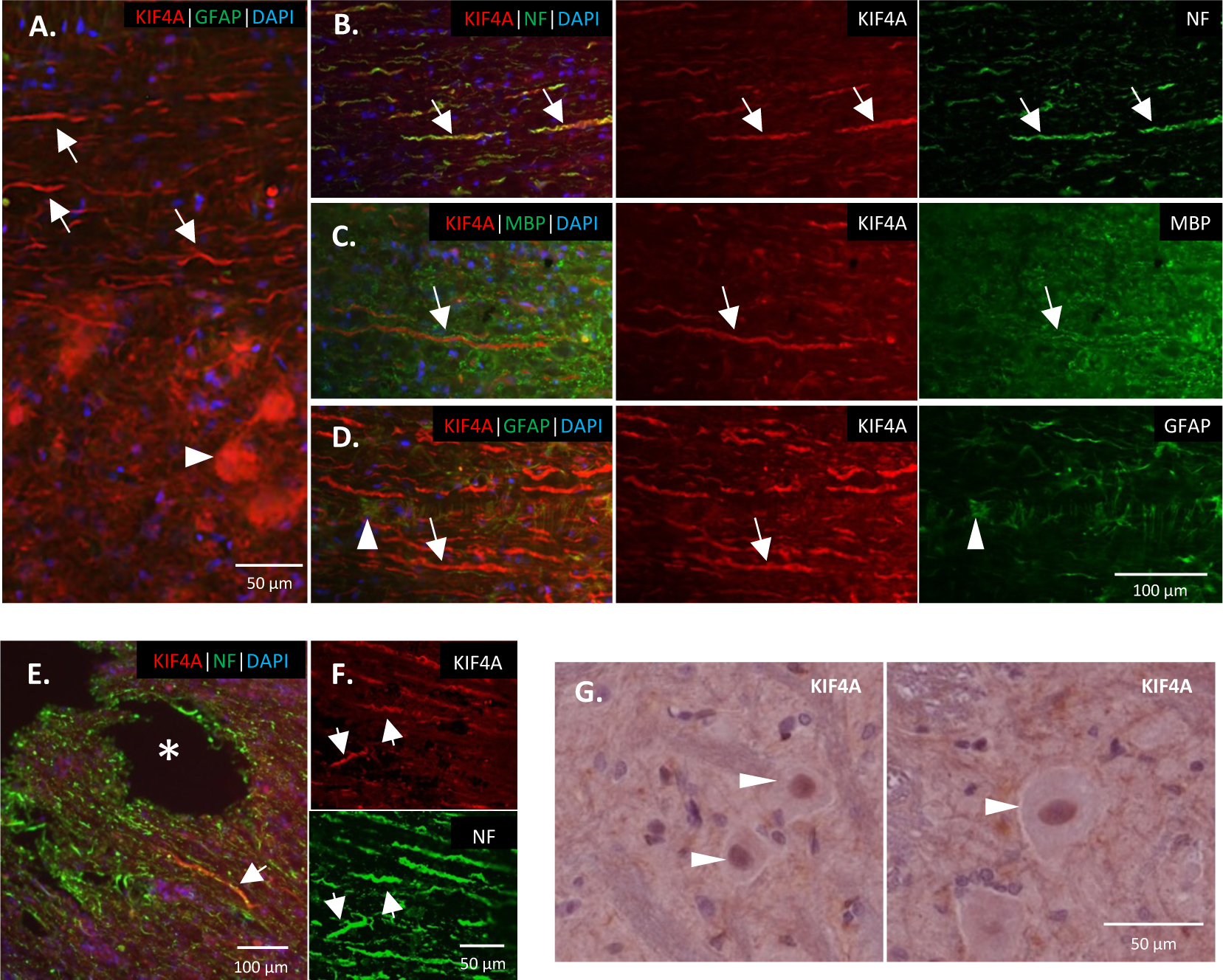
KIF4A immunostaining of the rat spinal cord. **A-D.** Fluorescence microphotographs of KIF4A (immunostained in red) in 50 µm sagittal cuts of an intact spinal cord; **E-F.** 50 µm sagittal cuts of a transected spinal cord. **A, B** and **F.** Co-labelling with neurofilaments (NF, in green), shows KIF4A/NF overlapping in axons (arrows). **C.** Co-labelling with the myelin marker MBP (green) showed a juxtaposed labelling of KIF4A and MBP, as in myelinated axons. **D.** Staining with GFAP, a glial marker, shows markedly different KIF4A-positive (arrows) and GFAP-positive (arrowheads) structures. **E.** Sections of a lesioned spinal cord, 3 days after lesion, where KIF4A still overlaps with NF (arrows); *, lesioned area. **G.** Colorimetric microphotographs of KIF4A expression in neuronal cells nuclei of paraffin-embedded spinal cord sections, 10 days after a peripheral nerve injury. Overall, KIF4A is present in both grey (**A** and **G**, arrowheads indicating cell bodies) and white matter (**A-F**, arrows indicating axons).

In sagittal sections of an injured spinal cord (**Figure 1E-F**), overlap of KIF4A with NF continued to be evident, indicating that this kinesin retains its axonal distribution after lesion. Likewise, KIF4A nuclear labelling of larger neuronal cell bodies of the spinal cord grey matter persists after lesioning of a downstream peripheral nerve (**Figure 1G**).

### KIF4A is present in both mature dorsal root ganglia neurons and peripheral axons

Subsequent analyses of KIF4A IHC labelling in the PNS demonstrated that this protein is located in both neuronal DRG cell bodies of mature uninjured rat dorsal root ganglia (**Figure 2**) and in axons of a sciatic nerve (**Figure 3**). Detailed analysis of DRG cross-sections reveals that KIF4A is expressed by different neuronal subtypes, as IB4-positive smaller DRG neurons and NF200-positive larger DRG neurons. Subcellularly, KIF4A is found in the cytoplasm of both IB4- and NF200-positive neurons but is also highly abundant in the nuclei of smaller and bigger DRG neurons (**Figure 2**, arrows). In addition, KIF4A is located in contact zones of neurons and surrounding satellite cells^51^, and may be also expressed by the latter (**Figure 2**, arrowheads).

**Figure 2.**
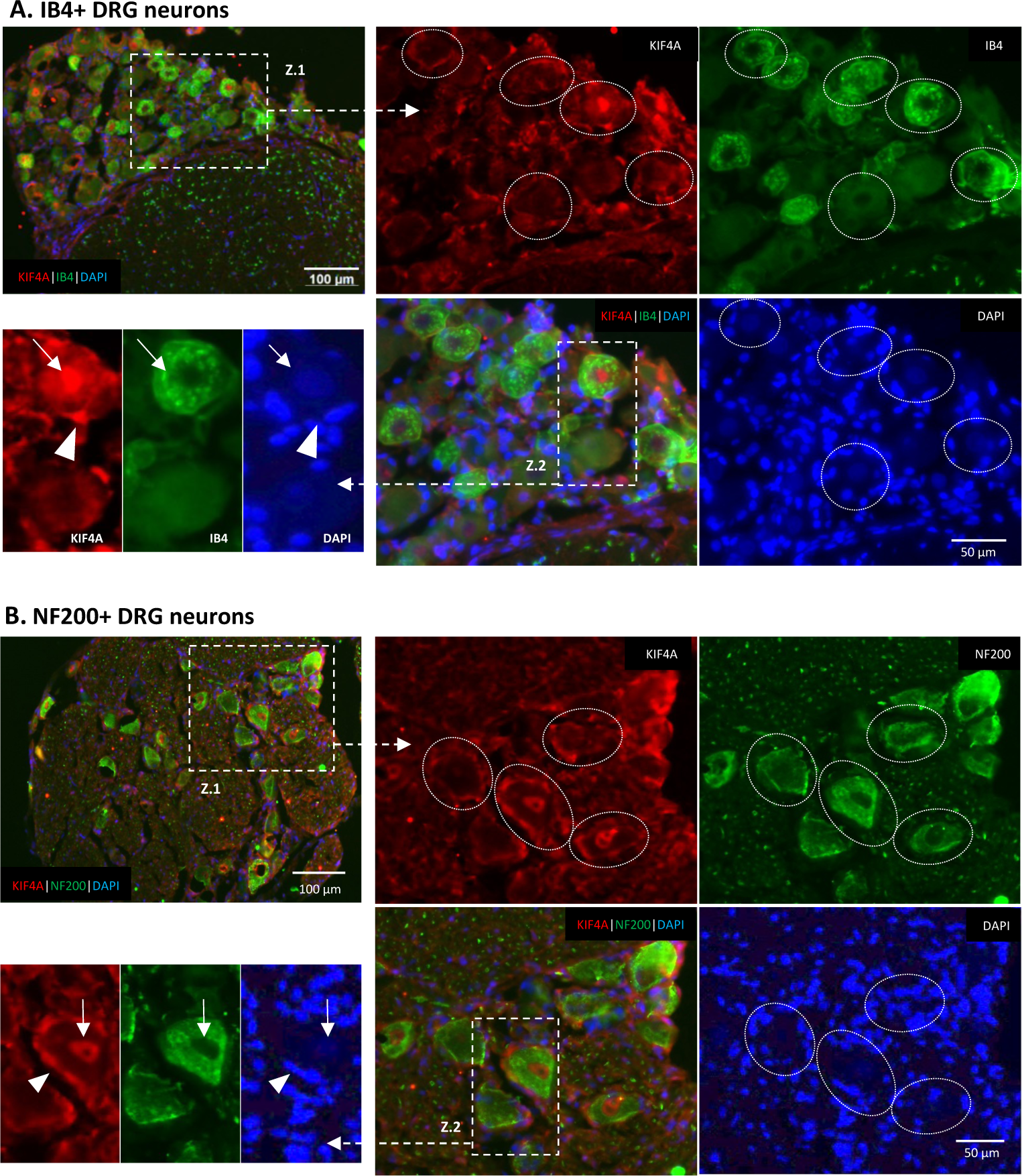
Fluorescence immunostaining of KIF4A distribution in cross sections of uninjured rat dorsal root ganglia (DRG). **A.** IB4 dye (in green) stains the smaller neurons in DRG. **B.** Neurofilament NF200 (in green) is marked in the larger DRG neurons. KIF4A (in red) stains some nuclei of both types of neurons (faint blue staining, zoomed pictures, arrows), and satellite cells surrounding the neurons (arrowheads). Z1-2, zoom 1 and zoom 2. Brightness in DAPI channel was intensified to reveal the less intense neuronal nuclei.

**Figure 3.**
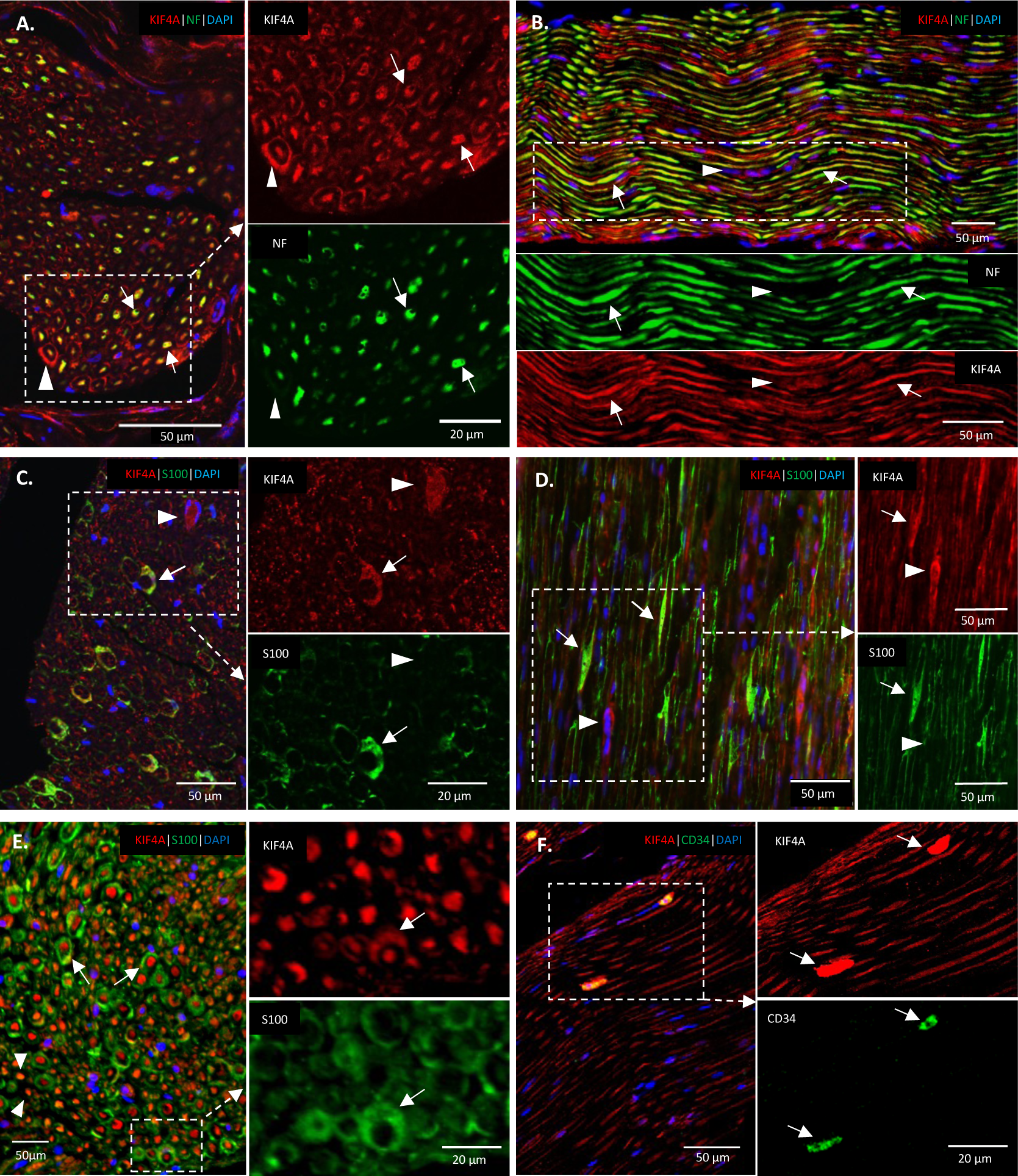
Fluorescence immunohistochemistry analyses of KIF4A protein expression in rat uninjured peripheral nerves. **A, C, and E.** Sciatic nerve cross-sections, and **B, D, and F.** longitudinal sections, of a rat sciatic nerve. **A** and **B.** Immunofluorescence analyses show that KIF4A (in red) co-labels with neurofilaments (NF) and is highly abundant in the nerve axons (arrows); KIF4A is also present in non-axonal structures (arrowheads). **C, D and F.** Co-labelling with the S100 marker identified most of the extra-axonal structures as Schwann cells (arrows), although there are also KIF4A-positive/S100-negative structures (arrowheads). **E.** KIF4A seems present in both myelinated (arrows) and unmyelinated (arrowheads) axons. **F.** KIF4A also co-labels with the endothelial marker CD34 (arrows).

In the intact rat sciatic nerve, double labelling of KIF4A with the NF axonal marker revealed a high number of tubular structures positive for both markers (**Figures 3A-B**, arrows), confirming that KIF4A is abundant in sciatic nerve axons. KIF4A appears to be present in both myelinated (**Figure 3E**, arrows) and unmyelinated axons (**Figure 3E**, arrowheads), although these latter are difficult to label and identify under light microscopy. Other cellular structures, whose nuclei can be seen on longitudinal sections and appear to surround axons are also KIF4A-positive but NF-negative (**Figures 3A-B**, arrowheads). Simultaneous labelling with an anti-S100 antibody confirmed that various of these KIF4A-positive structures were Schwann cells, where this kinesin appears to localize in the nucleus, cytoplasm, and plasma membrane (**Figure 3C-D** arrows). Additional KIF4A positive structures, neither labelled as axons nor as Schwann cells, were observed (**Figure 3C-D**, arrowheads). Double labelling with anti-CD34, a marker for hematopoietic stem/progenitor cells as well as for endothelial cells^52^, and with anti-neural/glial antigen 2 (NG2,) a perineurial and fibroblast-like cells marker^53^, revealed that KIF4A is expressed by both endoneurial fibroblast-like^54^ and endothelial cells, in both rat and human nerves (**Figure 3F** and **Supplementary Figure 1**, arrows).

Staining of KIF4A in an intact adult human nerve (**Figure 4**) confirmed the previous findings of clear axonal expression, visualized by positive dots at the core of fibers in nerve cross-sections (**Figure 4A** and **4C**, arrowheads), or as tubular structures in some longitudinal areas of a nerve section (**Figure 4B** and **4D,** arrows). Human Schwann cells also considerably express KIF4A (**Figures 4E-F**) in the same subcellular structures described above for rat (nucleus, cytoplasm and plasma membrane). Moreover, KIF4A seems to be present in both myelinated (**Figure 4F**, arrows) and unmyelinated axons (**Figure 4F**, arrowheads) in human nerves. Other unidentified KIF4A-positive cells were also observed in the human nerve (**Figure 4F**, asterisk).

**Figure 4.**
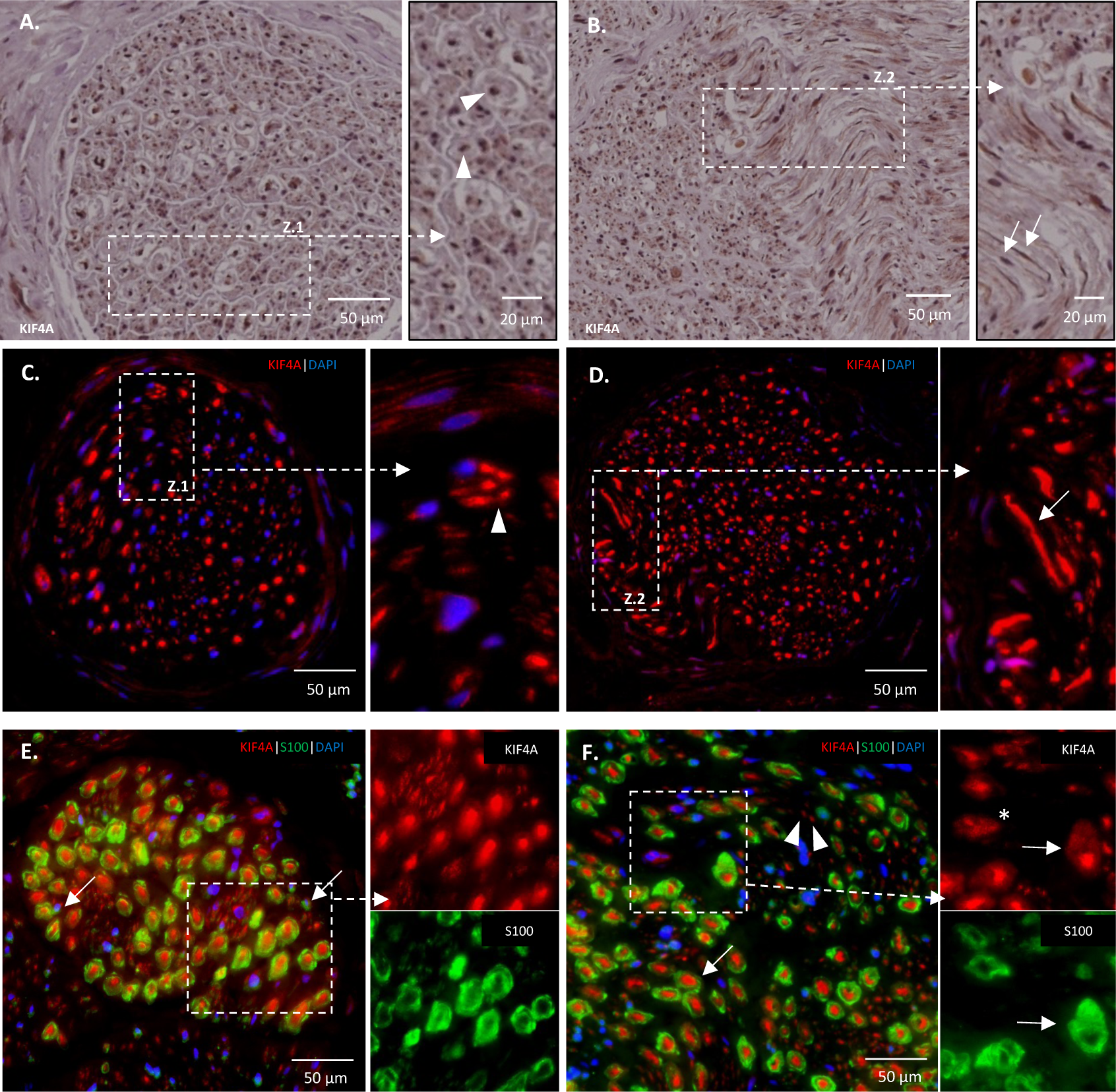
Fluorescence immunohistochemistry analyses of KIF4A protein expression in human uninjured peripheral nerves. **A-B.** KIF4A chromogenic and **C-D.** KIF4A fluorescence immunostainings, of cross-sections of a human craniofacial nerve. The same kinesin axonal pattern, previously observed in rat nerve, is shown. Axonal bundles of unmyelinated fibers (arrowhead in **A** and **C**), as well as typical longitudinal portions of myelinated axons (arrows in **B** and **D**) were also found in these transversal sections. **E-F.** KIF4A co-labelling with S100 detected most KIF4A-positive extra-axonal components as human Schwann cells (arrows in **E**). Positive axonal KIF4A detection was confirmed in myelinated (arrows in **F**) and unmyelinated (arrowhead in **F**) peripheral nerve fibers. However, some S100-negative cells also express KIF4A (asterisk in **F**).

### Injury-induced *Kif4a* expression in sensory DRG neurons

Considering the confirmed expression of KIF4A in adult CNS and PNS neurons, such as motor (**Figure 1**) and sensory DRG (**Figure 2**) neurons, further analyses focused on its possible modulation during regeneration. To verify whether *Kif4a* expression was altered following PNI, *Kif4a* mRNA levels in lesioned DRGs were analyzed at various days post-injury (dpi), by real-time PCR. Two experimental PNI models, with different severity and extent of subsequent regeneration, were used: (i) crush injury and (ii) transection of rat sciatic nerve with ligation. In the crush paradigm, mechanical compression results in a disruptive lesion of the axons and their surrounding myelin sheaths, leading to Wallerian degeneration processes distally to the lesion site. However, since the outer connective tissue layers (perineurium and epineurium) remain largely intact, regeneration tends to be complete. Nerve transection leads to a complete discontinuation of the whole nerve and of its surrounding connective tissue, and to subsequent axonal degeneration of the distal stump^14^. To impede successful regenerative ingrowth of proximal axons into the distal fragment, additional ligation (suture) was applied to the transected nerve stumps.

qPCR data of DRG neurons over time revealed lesion-induced *Kif4a* mRNA regulation (**Figure 5A**). Upon sciatic crush, *Kif4a* exhibits maximal up-regulation (∼2-fold, p-value <0.01) at 2 dpi and slowly declines back to basal levels of uninjured sham condition within 2-3 weeks. After transection, a similar expression profile was observed, with a slightly stronger initial up-regulation (∼2.7-fold, p-value <0.05) and a subsequent slow decline to basal levels.

**Figure 5.**
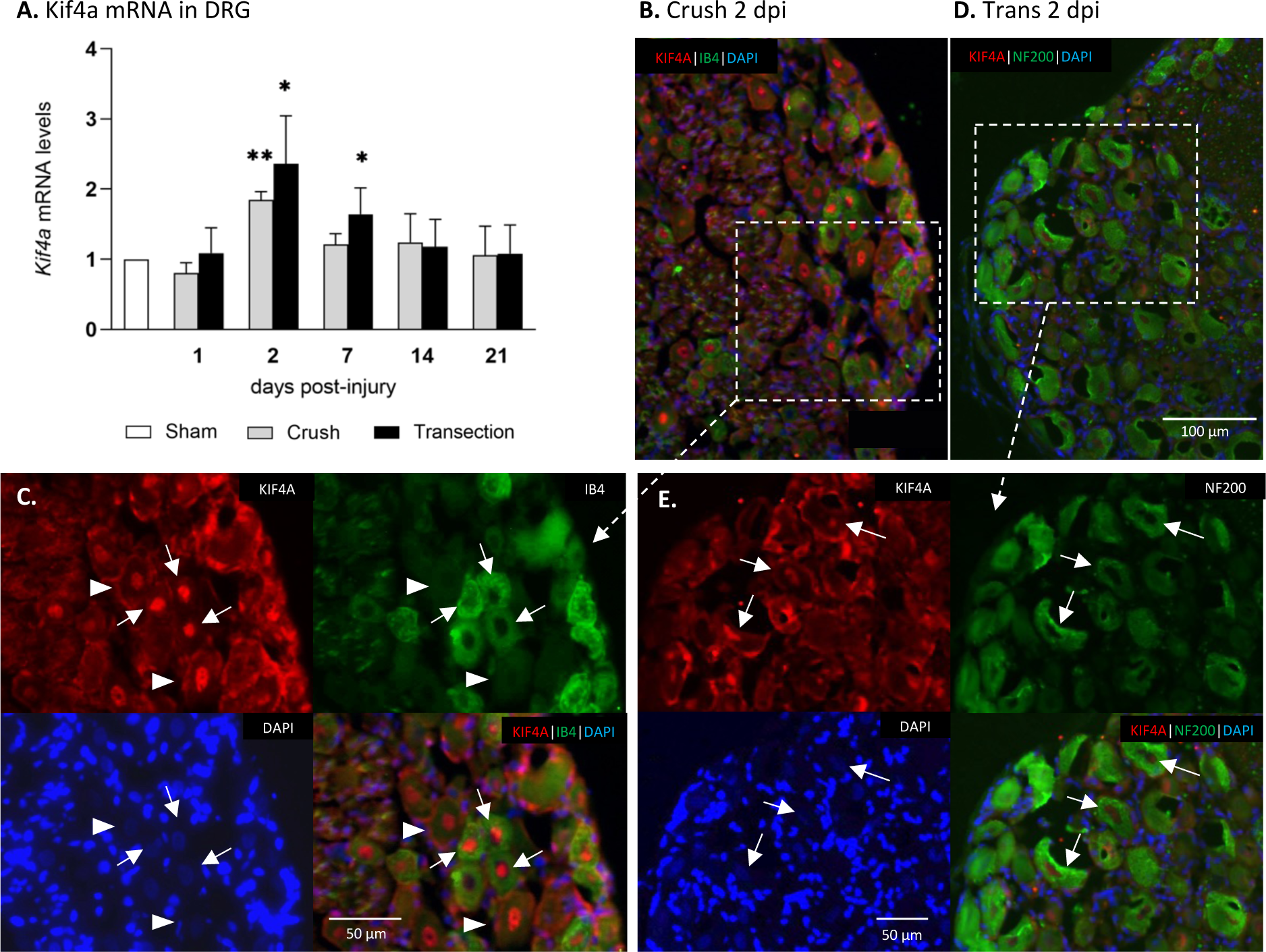
*Kif4a* expression and protein distribution in DRG neurons affected by peripheral nerve injury. **A.** Gene expression analysis of *Kif4a* mRNA levels in rat sensory DRG neurons at various time points following crush and transection lesions, by real-time PCR. Values are presented as mean±SD fold increases over mRNA levels of control sham-operated animals (taken as ‘1’); n=3; *, *p-*value <0.05; **, *p-*value <0.01, using unpaired t-test with Welch correction. **B-E.** Fluorescent microscopy analyses of KIF4A immunostaining in cross-sections of DRG at 2 days post-injury (dpi) via nerve crush (**B-C**) or transection with suture (**D-E**). Neurons were labelled with IB4 (**B-C**) or NF200 (**D-E**). In the crush condition, a high number of neurons is visible with abundant KIF4A staining in their nuclei (**C**, arrows for IB4-intense neurons; arrowheads for IB4-faint neurons). Some of these neurons are also visible in the transected condition (**E**, arrows).

Parallel immunohistochemical analysis of KIF4A in rat DRG injured neurons at 2 dpi revealed the same above stated KIF4A distribution in the neuronal cytoplasm and plasma membranes (and/or satellite cells) after both crush and transection lesions (**Figure 5B-E**). Nevertheless, KIF4A expression appears to be increased in the nuclei of both small IB4-positive neurons (arrows) and large unstained neurons (arrowheads) after crush injury, which correlates temporally with the activation of a variety of regenerative processes (**Figure 5B-C**, crushed condition). In contrast, in the transection with ligation paradigm, a rather subtle KIF4A increase is seen only in some neuronal nuclei (**Figure 5E**, arrows).

### *Kif4a* expression increases in the distal nerve stump upon sciatic nerve injury

Alterations in *Kif4a* expression were also monitored in the distal stump, a region of the nerve tissue that is particularly subjected to large alterations after traumatic injury.

An injury-induced temporary *Kif4a* up-regulation could be observed in the distal stumps of both crushed and transected sciatic nerves. *Kif4a* mRNA expression was significantly increased already at 2 dpi, reaching a maximum at 7 dpi in both lesion paradigms (∼12-13-fold increase over sham-operated animal levels; p-values <0.01 for transection and <0.05 for crush conditions) (**Figure 6A**). *Kif4a* levels subsequently declined over time back toward basal sham levels in both lesion paradigms, with crush levels remaining slightly above those of transection.

**Figure 6.**
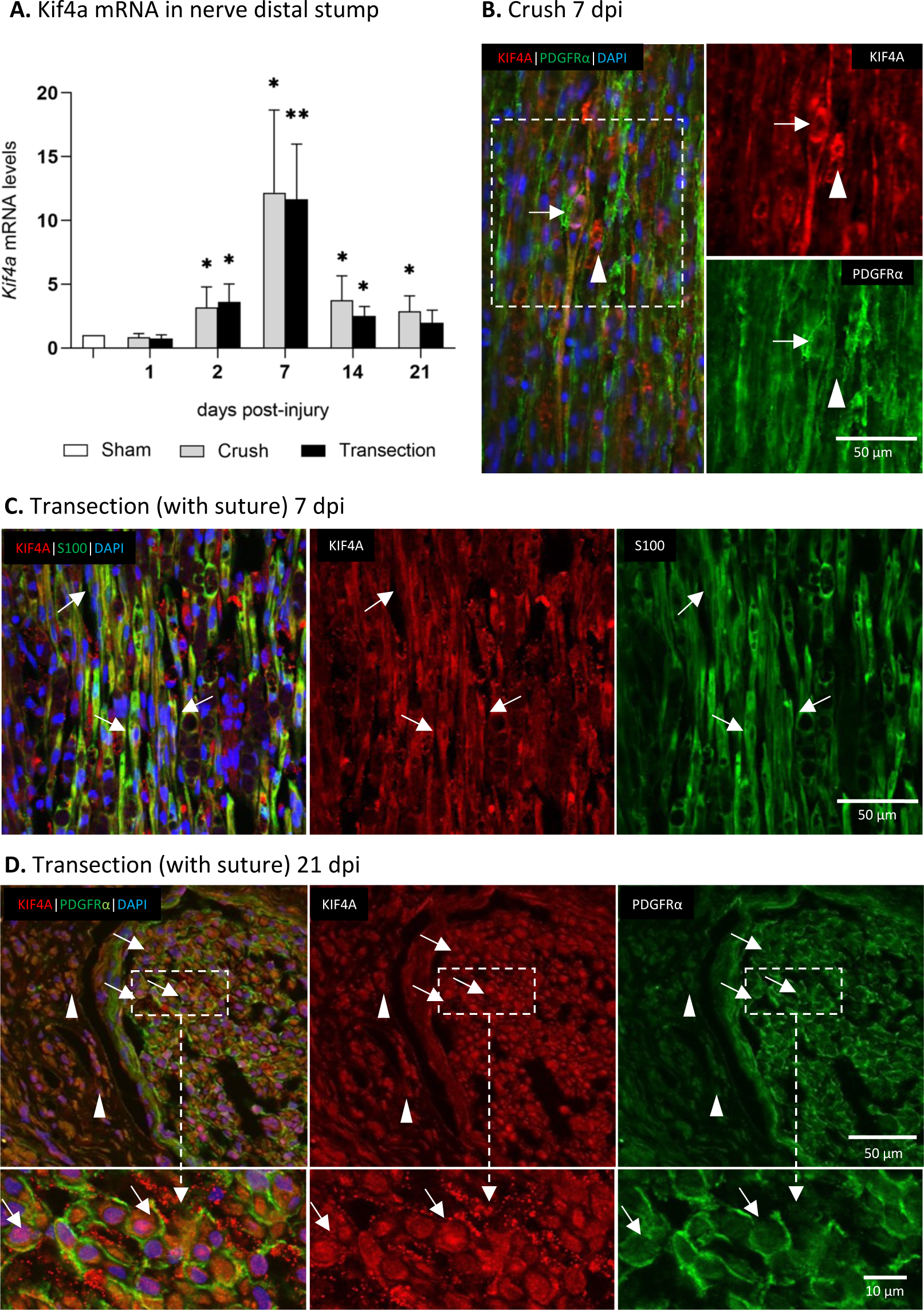
*Kif4a* up-regulation and protein distribution in a distal nerve stump with time post-injury. **A.** Gene expression analysis of *Kif4a* mRNA levels in the distal stump of a sciatic nerve at various time points following crush and transection lesions, by real-time PCR. Values are presented as mean±SD fold increases over mRNA levels of control sham-operated animals (taken as ‘1’); n=5; *, *p-*value <0.05; **, *p-*value <0.01, unpaired t-test with Welch correction. **B-D.** Fluorescence immunohistochemistry analyses of the lesion site of a (**B**) crushed sciatic nerve, longitudinal section, collected at 7 days post-injury (dpi); (**C-D**) transected and sutured sciatic nerve, longitudinal section collected at 7 dpi (**C**) and cross-section collected at 21 dpi (**D**), co-immunostained for KIF4A (in red) and for PDGFR-α (**B, D**, in green, staining glial cells) and for S100 (**C**, in green, staining Schwann cells). Arrows: cells double stained, confirming KIF4A to be up-regulated in Schwann and other supporting cells; arrowheads: cells/structures only stained by KIF4A.

Further IHC results indicate the histopathologic responses to both injuries in the distal stumps. Whereas processes of axonal degeneration, transdifferentiation and proliferation of Schwann cells, and regenerative axonal ingrowth, occur in the distal stump after crush injury, the latter is absent in the ligated transected distal stump^22,55^. Indeed, anti-neurofilament staining of transected distal stump sections at either 7 dpi or 21 dpi revealed complete axon degeneration (**Supplementary Figure 2A** and **B**). In contrast, stained nerve tissue of a crushed distal nerve fragment taken after 7 dpi adjacent to the lesion area, appeared relatively normal (uninjured) in sagittal sections (**Figure 6B** versus **Figure 3D**). Consistently, increased mRNA levels of octamer-binding factor 6 (Oct6), a marker for promyelinating Schwann cells^56^, were observed in the crushed distal stump at 7-21 dpi, indicating axonal induction of regenerative remyelination processes. Contrary, Oct6 mRNA levels remained very low in the transected and ligated fragment (**Supplementary Figure 2C**).

These data suggest that the distinct lesion-induced *Kif4a* upregulation observed in both crushed and transected distal nerve stumps occur in non-neuronal cells, such as distal Schwann cells, independent of the regenerative myelination process. Indeed, in immunohistochemical staining of the injured distal fragment, KIF4A protein signal was clearly detected in S100-positive Schwann cells (arrows in **Figure 6B-D**) labeled with S100 (**Figure 6C**). Complementary staining with the glial and endothelial marker PDGFRα (**Figures 6B** and **D**) also revealed clear similarities in KIF4A cellular expression. Additional, although less frequent, KIF4A-positive but PDGFRα-negative cells were observed, which could correspond to fibroblasts or other endoneurial cells (**Figures 6B** and **D**, arrowheads).

### KIF4A in newly formed axons and Schwann cells of a biomaterial-assisted regenerating nerve

KIF4A has been associated with anterograde transport of growing axons during development^39,40^. Given its axonal distribution in the adult uninjured nerve (**Figure 3**) and the observed early increase in *Kif4a* mRNA levels in regeneration-able injured sensory neurons (**Figure 5A**), an established nerve conduit implant was used for neuronal regeneration supplementary studies. The NeuraGen® Nerve Guide Collagen Type I Conduit is an implant for bridging peripheral nerve discontinuities that provides a protective environment for peripheral nerve repair after injury^10^. This conduit was used to bridge the proximal and distal stumps of a transected rat sciatic nerve and to promote axonal outgrowth of proximal axons^10^ (**Figure 7A**). Staining of the proximal axons grown distally through the conduit at 10 dpi showed a distinct fibrous GAP43 distribution of regenerating axons, with a focus in the left half of the distal cross-section (**Figure 7B**). KIF4A staining in an immediately adjacent section showed a clearly corresponding distribution (**Figure 7C**). Higher magnification images suggest that KIF4A is expressed not only in the ingrown axons but also in their neighboring cells, presumably Schwann cells (**Figure 7C**, magnification). To confirm this assumption, co-staining with KIF4A and the Schwann cell marker S100 was performed on nerve cross sections that were adjacent to the above sections (**Figure 7D**). As expected, many cells in the distal intraluminal regenerating nerve tissue were positive for both S100 and KIF4A, confirming KIF4A abundant expression in regeneration-associated repair Schwann cells.

**Figure 7.**
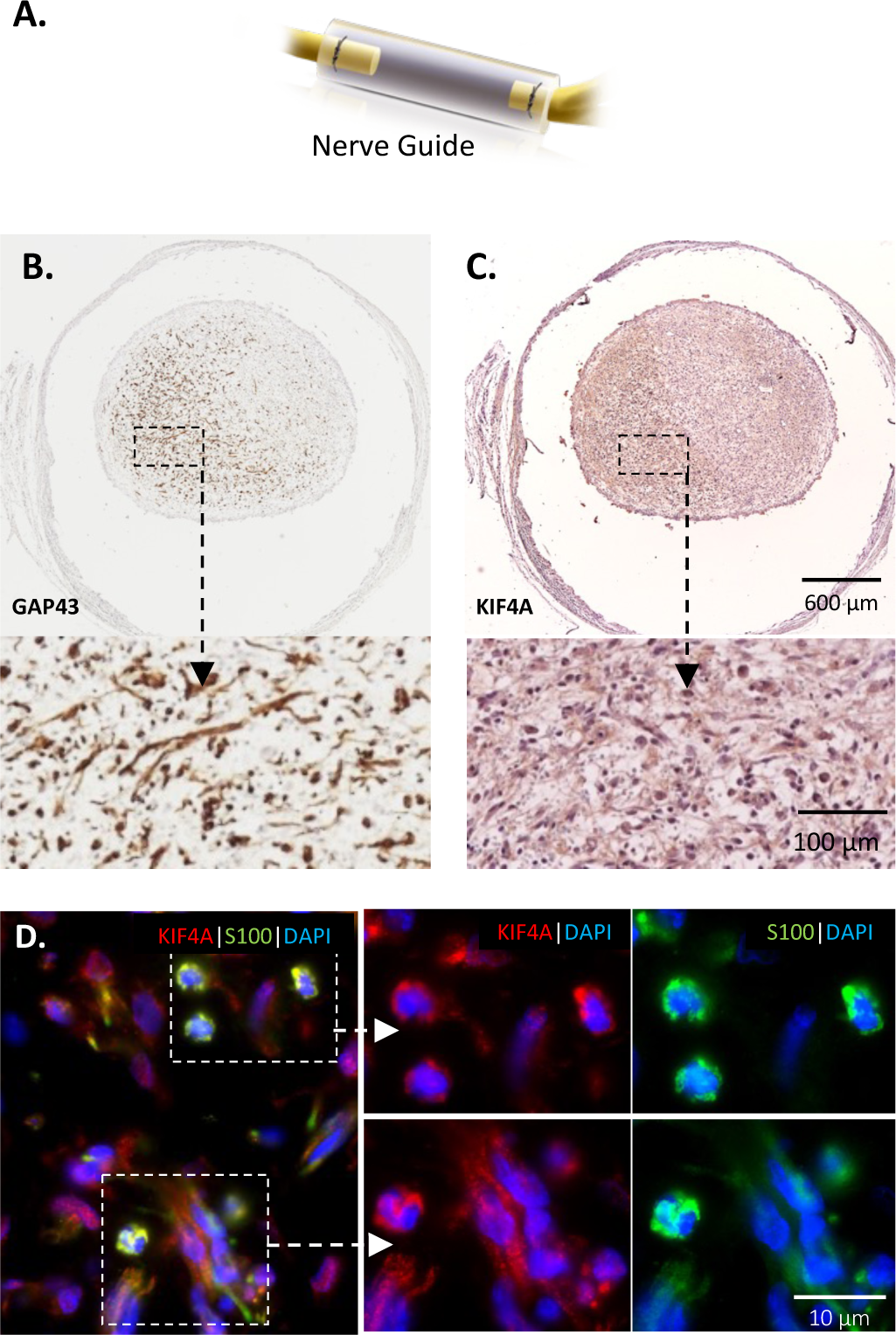
KIF4A immunostaining in rat regenerating transected sciatic nerve fascicles upon conduit application. **A.** Schematic image of the conduit “Neuragen® Nerve Guide” (adapted from Integra LifeSciences Corporation). **B-D.** Microscopy analyses of KIF4A distribution at 10 days post-injury in a transected and repaired sciatic nerve using the NeuraGen conduit. Immunohistochemical detection of GAP43 (**B**) and KIF4A (**C**) in adjacent sections, inside the conduit, showing a diffuse positive reaction in all the regenerating nerve tissue, but particularly intense in the left half of the regenerating nerve. **D.** Immunofluorescence analysis of KIF4A and S100 signals in the conduit-supported regenerating nerve evidences a high co-labelling of both staining that confirms that KIF4A is expressed in repair supporting Schwann cells.

### *Kif4a* is induced in proliferative repair Schwann cells while *Kif4a* silencing hinders proliferation

As shown above, damage to a peripheral nerve whether by crush or transection, causes a temporally parallel increase in *Kif4a* expression in both sensory DRG neurons and Schwann cells of the distal nerve stump. During first days after injury, myelin and non-myelin (Remak) Schwann cells distal to the nerve injury undergo a large-scale change in function, from maintaining the axonal sheath and myelin to supporting regeneration^14,57–59^. This switch to a ‘repair’ phenotype is accompanied by strong cellular proliferation^60^. Using primary Schwann cell cultures, which mimic to some extent the immature-like status of repair Schwann cells^56^, we further tested if the proliferation activity of cultured Schwann cells correlated with *Kif4a* expression. To this aim, the *Kif4a* mRNA levels in Schwann cells in proliferating sub-confluent cultures was compared to the one in non-proliferating density-arrested of confluent cultures. Proliferating Schwann cells grew as subconfluent layers reaching a maximum cell density of 40-50%, and dense confluent layers of Schwann cells were maintain for 24h after reaching ∼100% cell density, for cell contact-inhibited proliferation. Confirming our hypothesis, *Kif4a* mRNA levels were significantly (∼6-fold) higher in proliferating Schwann cells than in arrested cells (**Figure 8A**, qPCR graph). To better characterize these cultures, a parallel analysis was performed on the levels of two transcription factors, *Krox20* (EGR2) and *Oct6* (SCIP/Tst1), considered necessary for Schwann cells transition from a promyelinating to a myelinating stage during *in vivo* development. Both factors were highly decreased in the more proliferative condition and increased in the cultured arrested Schwann cells (**Supplementary Figure 4**), indicating a more differentiated state in this last condition. Altogether, these suggest that *Kif4a* expression peaks in the more immature proliferating Schwann cells.

**Figure 8.**
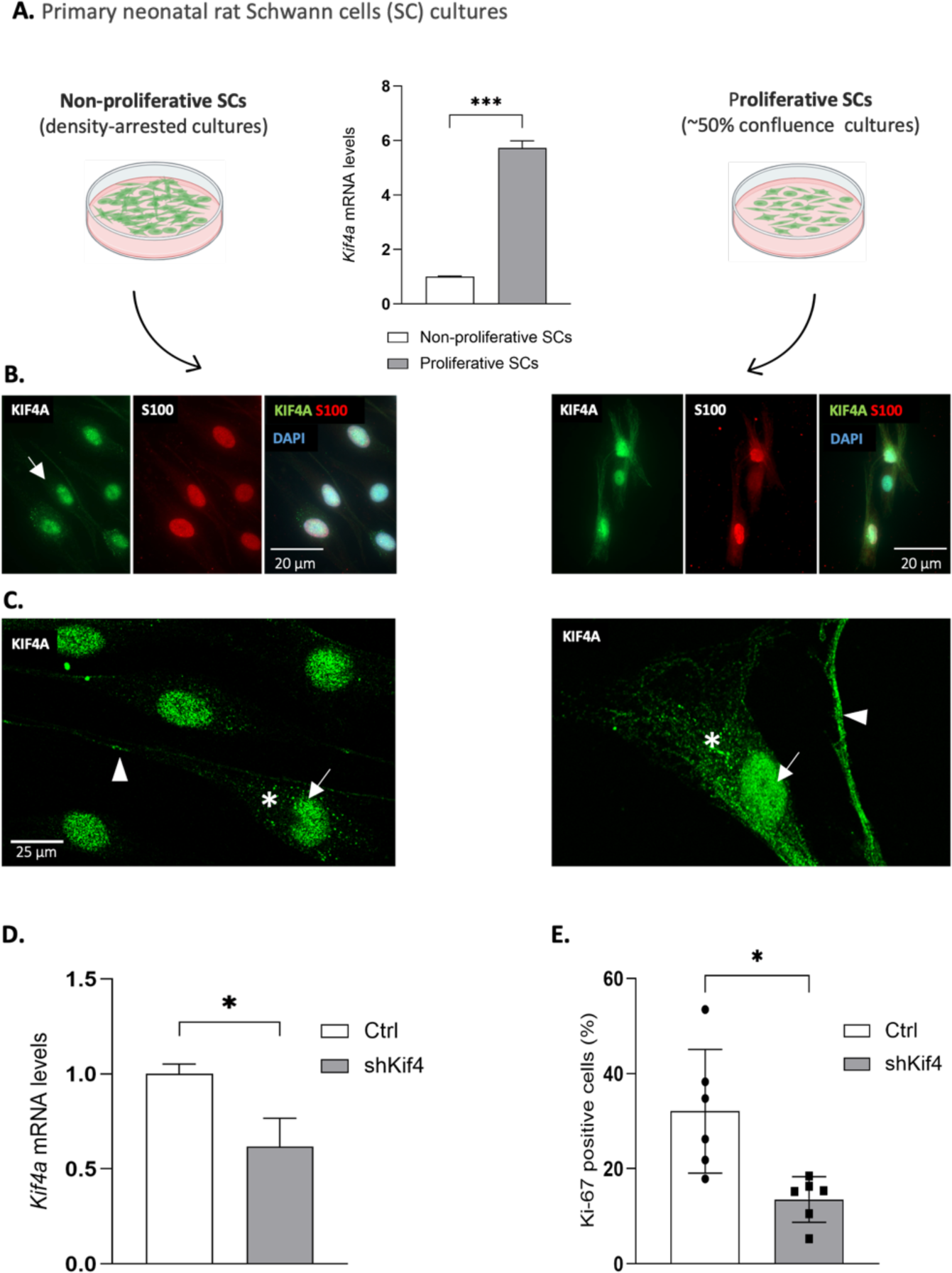
*Kif4a* expression in proliferative and non-proliferative Schwann cells and *Kif4a* role in Schwann cells proliferation. **A.** Schematic representation of the rat cultured Schwann cells (SCs) models used (confluent non-proliferative and sub-confluent proliferative), and graphic analysis of *Kif4a* mRNA levels in both proliferation-state models, quantified by real-time PCR; n=3. **B.** Fluorescence microphotographs of KIF4 (in green) and S100 (in red) staining in SCs in both proliferation-state models. **C.** Higher magnification confocal microphotographs of cells in both conditions, highlighting KIF4A subcellular distribution: high nuclear abundancy (arrows), and presence in cytoplasmic structures (asterisks) and cellular projections (arrowheads), with the two latter being particularly visible in proliferative cells (microphotographs taken with equal settings). **D** and **E.** *Kif4a* role in SC proliferation. **D.** *Kif4a* mRNA levels assessed by qRT-PCR upon 48h of transfection with an anti-*Kif4a* shRNA GFP-expressing plasmid; cells transfected with the empty plasmid (only GFP-expressing) were used as controls. Results expressed as mean±SD; n=3. **E.** The percentage of proliferating Ki-67-positive cells (immunostained in red) was scored in green fluorescing cells (expressing shKif4 and GFP, or only GFP); bar in the dispersion graph represents the mean±SD of n=6, 200-500 cells scored per ‘n’. *, *p-value* <0.05; ***, *p-value* <0.001, using unpaired t-test with Welch correction.

Complementary immunocytochemical staining shows that in both conditions KIF4A was predominantly located at the Schwann cells nuclei (**Figure 8B-C**, arrows), as previously observed in Schwann cells in the transected nerve distal stump (**Figure 6C**) and in the conduit-assisted regenerating nerve (**Figure 7D**). However, the KIF4A nuclear staining of density-arrested Schwann cells appeared to be less intense than in proliferative cells and, in these latter, KIF4A was also localized in several cytoplasmic structures apparently aligned with microtubule filaments (asterisks), as well as in cellular projections (arrowheads) (**Figure 8C**, confocal higher magnifications). To better characterize these cultures, a parallel analysis was performed on the levels of two transcription factors, *Krox20* (EGR2) and *Oct6* (SCIP/Tst1), considered necessary for Schwann cells transition from a promyelinating to a myelinating stage during *in vivo* development. Both factors were highly decreased in the more proliferative condition and increased in the cultured arrested Schwann cells (**Supplementary Figure 4**), indicating a more differentiated state in this last condition. Altogether, these results indicate that KIF4A expression peaks in the more immature proliferating Schwann cells, where it localizes not only in the nucleus but also in cytoplasmic structures.

To confirm a putative role of *Kif4a* in Schwann cells’ proliferation, endogenous *Kif4a* expression was suppressed in Schwann cells grown under the proliferative culture conditions, using a specific anti-*Kif4a* shRNA plasmid, also GFP expressing (**Figure 8D**, **Supplementary Figure 5**). Subsequent scoring of the percentage of cell division-active Ki-67-positive cells in the GFP-expressing transfected cells population, revealed a decrease in the number of proliferating cells by approximately 50% due to shKif4a-mediated suppression (**Figure 8E**). This result confirms our hypothesis of a causal relationship between *Kif4a* expression and the proliferation activity of Schwann cells.

## DISCUSSION

Development and regeneration of the nervous system depend on motor proteins of the kinesin superfamily (KIFs), to e.g. perform anterograde transport to the microtubules’ plus-ends and promote growth cone progression^33,61,62^. KIF4A is a microtubule-binding protein associated with neuronal development disorders and, until now, only reported to be expressed during the embryonic phase. Its known functions in the developing nervous system were mainly to transport cargoes from the cell body of e.g. embryonic cortical and DRG neurons to the axonal tip, promoting axonal growth.

Our study demonstrates for the first time that KIF4A is also widely expressed in the central and peripheral adult nervous systems of both rats and humans. It was detected in the spinal cord’s white and grey matter (**Figure 1**), in the cell bodies of adult rat DRG neurons (**Figure 2**), and in axonal tracts of rat and human peripheral nerves (**Figures 3 and 4**). Consistent with its suggested roles as an axonal kinesin and as a chromokinesin, KIF4A is not only expressed in axonal tracts but is also abundant in the nuclei of neurons and other neural PNS cells. Indeed, KIF4A was detected to a high extent in Schwann cells (further discussed below), and also in some S100- or PDGFRα-negative cells that positively stained for CD34 or NG2. These exhibited a morphology similar to the one of endoneurial fibroblast-like cells (EFLCs), known to maintain nerve structure and promote axonal regeneration^63–65^.

### KIF4A in regenerating injured neurons

Evaluation of lesion-induced *Kif4a* expression was performed using two rodent PNI models of nerve injury and regeneration^66^: (i) the crushed sciatic nerve, a commonly used model for spontaneous regeneration, and (ii) the transection model with nerve stumps ligation, that prevents efficient ingrowth of regenerating axons into the distal stump^14^. RAGs temporal up-regulated in crushed neurons undergoing regeneration are often also up-regulated in neurons whose axons were cut and sutured, sometimes even at higher amplitudes and/or over longer time periods^14,26^, suggesting a robust and persistent attempt to promote regeneration via key pathways triggered by these RAGs. Accordingly, *Kif4a* mRNA levels were up-regulated in DRGs of both PNI conditions, early after injury (2 dpi), and the upregulation appeared to be slightly longer (7 dpi) under transection conditions (**Figure 5A**). Given the marked KIF4A staining in neuronal DRG nuclei/cell bodies, the observed regulation is likely attributed to the injured sensory neurons in particular, although involvement of neighboring satellite cells and Schwann cells cannot be excluded at this time^67^. Importantly, no other study so far has reported KIF4 neuronal up-regulation following injury, possibly due to the use of detection techniques of lower sensitivity, or to the analytical timepoints chosen (e.g. Northern blotting analysis of various KIFs in adult mouse DRGs at 7 and 14 dpi^37^). Further, under conditions in which regeneration of a severed rat sciatic nerve was enhanced by implantation of a conduit^68^, KIF4A was highly abundant in axonal tracts also stained for GAP-43 (**Figure 7B-C**). This established RAG is strongly up-regulated during neuronal regeneration and correlates with enhanced regenerative capacity^26,42^.

The results thus support our hypothesis that lesion-induced KIF4A upregulation reflects its dual neuronal regenerative role, both in the cell body and in the distal tips of regrowing axons. As part of the neuroregenerative response, a number of genes are known to revert to an embryonic and/or postnatal expression pattern after injury^14,69^. Accordingly, the lesion-induced *Kif4a* increased expression observed in our study mimics that of its neuronal development, and suggests the reestablishment of KIF4A functional tasks reported in developing neurons.

A main KIF4A role in embryonic neurons is the anterograde transport of proteins, ribosomes, and ribosomal components^40^ critical for axonal expansion in both developing and regenerative contexts. Anterograde transport, for example, is essential to satisfy the increased protein demand at the injury site and/or regrowing growth cone, especially for lengthier peripheral axons whose tips are quite distant from the cell body^61^. One of KIF4A cargoes in cultured cortical neurons is the widely known cell surface receptor β1-integrin^38^ that, among other roles, connects the actin cytoskeleton to the extracellular matrix (ECM) and is involved in cell adhesion and migration processes, and neurite growth^70^. In peripheral nerves and sensory axons β1-integrin is associated with neuronal polarization^71^and axonal outgrowth^72^. Noteworthy, β1-integrin (as KIF4A) levels decrease in mature central neurons^38,71^ and this decline is considered to be a main contributor for the remarkable loss of the adult CNS neurons regenerative capacity^71^. Importantly, exogenous KIF4A transfection resulted in apoptosis of cultured cortical neurons instead of increasing β1-integrin transport^38^, highlighting the crucial need to tightly control KIF4A levels. Another important KIF4A cargo transported into the axonal growth cones is L1, which can promote neurite elongation in embryonic cultured neurons^39,41^ and supports neuronal regeneration in both adult CNS and PNS^26,42^. All these facts strongly suggest a relevant function for KIF4A in neuronal intrinsic regeneration processes, although further studies are needed to investigate its precise role and mechanisms of action.

Aside its axonal transport role, KIF4A up-regulation in axotomized DRGs may also be related to a nuclear-associated function. Although histochemistry is not a reliable quantitative method, our immunostaining results suggest that nuclear KIF4A signaling appears enhanced in injured DRG neurons (**Figure 5B-C**). Previously, KIF4A has been associated with spindle stability during the cell cycle, apoptosis through PARP binding, and DNA repair through binding to chromosomes, all roles that may justify its neuronal upregulation and nuclear localization under both PNI conditions^62^. Activation of the cell cycle of nonmitotic cells such as neurons, and initiation of apoptosis (as e.g. the observed in the more severe transected injury model), are known biological processes after nerve injury^73^ in which KIF4A may be involved.

### KIF4A in pro-regenerative repair Schwann cells

In parallel with a presumed KIF4A neuronal role in promoting axonal regrowth, this study offers evidence, for the first time to our knowledge, of an indirect role of KIF4A in promoting regeneration of peripheral nerves by promoting proliferation of (repair) Schwann cells at the distal stump. The regeneration-promoting microenvironment of peripheral nerves is largely formed by non-neuronal neighboring cells such as Schwann cells, invading macrophages and EFLCs, as well as by the supporting ECM, which substantially stimulate neuronal regeneration processes and control axonal regrowth^21,74^. After nerve injury, Schwann cells dedifferentiate into an immature and proliferative state, and engage into an alternative differentiation program that culminates in a pro-regenerative phenotype^22^. These newly converted ‘repair’ Schwann cells will secrete neurotrophic signals and form the bands of Büngner, the key guidance cues in this complex regeneration process^16,22,75,76^.

We here observed a robust induction of *Kif4a* mRNA levels in both crushed and transected distal nerves, occurring over almost the entire period examined but starting as early as 2 dpi and with a clear peak at 7 dpi (**Figure 6A**). Our immunostaining of intact (**Figures 3 and 4**), injured and regenerating (**Figures 6 and 7**) neural tissues demonstrate KIF4A clear expression by Schwann cells (mainly) and other supporting cells. Further, considering that massive axonal degeneration occurs in the transected and ligated distal nerves, the observed ∼11-fold higher *Kif4a* expression cannot be of neuronal/axonal origin. Thus, the strong injury-induced *Kif4a* up-regulation in injured distal nerves can only be due to endoneurial cells, mainly Schwann cells, which represent the major cell group in the transected distal stump^60^. Accordingly, a search for *Kif4* in the supplementary transcriptomic datasets of a transected mouse sciatic nerve, by Arthur-Farraj and colleagues (2017)^60^, revealed an injury-induced ∼15.5-fold up-regulation in *Kif4a* mRNA levels at 7 dpi, similar to our data.

At this time point (7 dpi), the injury-induced myelin degradation, associated with Wallerian degeneration, is already well advanced. Schwann cells have lost their myelinating phenotype due to the loss of axonal contact and will or have already adopted a proliferative “repair” phenotype that is essential for successful nerve regeneration^60,77,78^. Our data shows that *Kif4a* regulation (**Figure 6a**), unlike that of *Oct6*, is independent of Schwann cells’ acquisition of a promyelinating phenotype by regenerative re-established axonal contact (only occurring in the crush condition; **Supplementary Figure 2**). Hence, KIF4A should act earlier, in more immature cells, and its chromokinesin functions in mitosis and spindle stabilization could be particularly relevant to the transdifferentiation and proliferation of Schwann cells in the distal stump, to expand their population for the formation of the regeneration-promoting Büngner’s bands. A role for KIFA4 in immature Schwann cells was already suggested by the data from rat Schwann cells cultures at different confluences, where proliferating cells expressed ∼5-6 times more *Kif4a* mRNA than growth-arrested ones (**Figure 8A**). This model was used since *in vitro* cultures of Schwann cells mimic to some extent the immature phenotypes of *in vivo* developing undifferentiated, and of adult dedifferentiated Schwann cells after nerve injury^56^. Further confirmation was provided by the significant ∼50% reduction in Schwann cells’ proliferation rate in cells where KIF4A expression was knock-down by anti-Kif4a shRNA means. These results confirm a KIF4A role in the proliferation of immature Schwann cells, and strengthen our assumption that injury-induced *Kif4a* expression in the distal stump is crucial for Schwann cells transition from a mitotically inactive mature myelinating phenotype into a proliferative repair phenotype^79^.

Considering that KIF4A is almost exclusively nuclear located in mitotically inactive Schwann cells while it is distributed in the nuclei and in cytoplasmic dots apparently microtubule-aligned in proliferative cells (**Figure 8B**), an extra-nuclear role is also suspected for KIF4A in repairing Schwann cells, such as transporting L1 to the surface, for mediating axon-Schwann cell contact. To gain more insight into possible KIF4A mechanisms of action in repair Schwann cells, we searched for known KIF4 interactors in *Mus musculus* [**Supplementary Figure 3A**, total KIF4 protein-protein interaction (PPI) network] within a list of differentially expressed genes (DEGs) at the injured distal nerve fragment at 7 dpi (Farraj’s work)^60^. A total of 32 *Kif4* interactors were found (plus *Kif4* itself): 23 upregulated and 9 downregulated (**Supplementary Figure 3B**). These included 5 MHC class II antigen presentation proteins of the Ig superfamily (including the KIF4A cargoes L1 and integrins), 17 kinesins and 1 dynein. Based on the PPI network obtained, KIF4A appears to be functionally involved in several cellular processes (at 7 dpi), including mitotic nuclear division, regulation of microtubule cytoskeleton organization, Golgi-to-ER retrograde traffic, trans-Golgi Network Vesicle Budding, transport along microtubule (**Supplementary Figure 3C-D**). The KIF4A cytoplasmic pools observed in Schwann cells (*in vivo* and *in vitro*) and in regrowing axons, suggest a microtubule-based transport function where the transported cargoes are presumably regeneration-promoting and/or involved in neuron-glial contact. The KIF4A cargoes β1-integrin and L1^39^ are known to be present in myelinating Schwann cells and in regenerating axons^80,81^, and to play important roles in neural regeneration^82–84^ and neural cell-cell trans-dimerization^85^. Hence, we hypothesize that KIF4A plays a dual role at the distal nerve, probably in a time subsequent manner: (i) promotion of the proliferation of transdifferentiated repair Schwann cells; (ii) promotion of the Schwann cells-regenerating axon contact, involving the essential transport of L1 and β 1-integrin to the cell surface^86^. This critical cell-cell contact is a prerequisite for the initiation of repair Schwann cells re-differentiation/maturation and axonal re-myelination^87^.

Finally, KIF4A is also expressed by additional cells of the injured distal nerve (**Supplementary Figure 1**), which are known to be important for axon regrowth following nerve injury^64^. For example, the CD34/NG2-positive EFLCs^54^ are observed in the proximal nerve stump as early as 1 dpi, and can assist Schwann cells during nerve regeneration by serving as a support for the bands of Büngner^63,88^.

## CONCLUSION

We here first show that the motor protein KIF4A, whose deregulation is associated with various neurodevelopmental pathologies, is expressed in adult CNS and PNS tissues. After PNI, KIF4A is not only induced in the cell bodies of axotomized neurons but is also strongly upregulated in non-neuronal cells of the injured nerve, which are involved in debris clearance and support axonal regeneration. On the one hand, in DRG neurons KIF4A can promote axonal transport of organelles and molecules important for axonal regrowth and is also recruited to the nucleus, where it can help overcome DNA and other damage and promote genomic stability ^89^. On the other hand, KIF4A is regulated in cells of the distal stump that promote regeneration, particularly in transdifferentiated Schwann cells that acquired a more immature and repair-relevant phenotype, where KIF4A is elevated to promote their proliferation and potentially axonal transport of relevant cargoes. These cells form Büngner’s bands, which are essential for axon regrowth and guidance. This dual role places KIF4A as an attractive target for regeneration therapies for PNI and SCI. Indeed, our observation of KIF4A expression in the adult spinal cord, and findings that *Kif4a* is significantly upregulated in *Xenopus laevis* at 2 days post-SCI but only in the regenerating tadpole, and not in the non-spinal cord regenerative froglet^29^, highly suggests a role for KIF4A in spinal cord repair. Considering that Schwann cells have considerable good efficacy rates in SCI cell therapy trials^90^, future combined genetic and cellular therapy, such as enhanced expression of KIF4A in implanted Schwann cells, may contribute to treat SCI. Nevertheless, further studies are needed to verify e.g. whether *Kif4a* absence leads to a significant loss of repair Schwann cells *in vivo*, which may subsequently result in failure of nerve repair and/or remyelination in the peripheral nerve. Clarification of the neuronal and Schwann cells KIF4A mechanisms to promote axonal regeneration after nerve injury may open new approaches for gene and cell therapy to treat CNS and PNS injuries to enhance neural regeneration.

## MATERIALS AND METHODS

### SCI and PNI animal models

Nerve injury experiments were conducted in adult Wistar rats (200-240 g) in compliance with a) the German Animal Protection Law and were performed with the permission of the State Office for Nature, Environment and Consumer Protection North Rhine-Westphalia (LANUV NRW, Recklinghausen, Germany) and b) the ethical committee of the University of Granada, Spain, following the European Union and Spanish Government guidelines for animal experimentation (EU Directive No. 63/2010, RD 53/2013), respectively.

For SCI experiments, rats were anesthetized with Isoflurane (Forene, Abbott; 2–3% in O_2_ and N_2_O at a ratio of 1:2). Following laminectomy at T8/9 and opening of the dura mater via a longitudinal cut, the spinal cord was slightly lifted and completely transected using micro-scissors. Spinal cords were collected 3 days after sham operation or transection.

PNI (sciatic nerve injury) experiments in anesthetized rats were prepared by a small skin incision over the gluteal region. The ischiocrural musculature was carefully spread to expose the sciatic nerve and nerve lesions (crush or transection) were bilaterally performed at the mid-thigh level. Severe axonotmetic crush injury – leading to axonal disruption but preserving the connective tissue – was performed using Dumont #7 jewelry forceps (2x15 sec)^14,66^. For neurotmetic transection injury, axons and connective tissue were fully cut. To ensure chronic denervation and degeneration of distal targets, the proximal stump of transected nerves was additionally ligated using 6/0 suture silk. Distal and proximal nerve fragments, and corresponding dorsal root ganglia (DRG, L4-L6), were collected at 1, 2, 7, 14 and 21 days after sham operation or nerve injury, respectively. Tissue samples were rapidly frozen after dissection, and stored at −30°C.

For nerve repair experiments, surgical procedures were performed upon general anesthesia by intraperitoneal injection of a mixture of acepromazine (Calmo-Neosan®, 0.001 mg per g of weight of the animal), ketamine (Imalgene 1000®, 0.15 mg per g of weight), and atropine (0.05 μg of per g of weight). When rats were anesthetized, a 10 mm segment was removed from the left sciatic nerve and subsequently repaired with NeuraGen® collagen type I conduits of 1.5 cm length and 0.2 cm internal diameter (Integra® Life Sciences Corp., Plainsboro, NJ, United States) following previously described protocols^91,92^. After surgery and wound closure, all rats were kept in individual cages and received metamizole in the drinking water as analgesia. Animals were kept for 10 days before euthanasia by an anesthesia overdose followed by intracardiac perfusion. First, we perfused 500 mL of physiological saline solution (to remove the blood from the tissues) and then 500 mL of 10% buffered formalin for the complete fixation of the animal. The grafted conduits and the perfused vertebral column were immediately harvested and post-fixed in 10% buffered formalin for 24 hrs. For the spinal cord tissues, the lumbar-sacral region (T13-L5) was carefully dissected and post-fixed for another 24 h (total fixation of 48 h).

Healthy sensitive human nerves were donated by patients who were subjected to reconstructive craniofacial surgeries, after obtaining their written informed consent. These tissue samples were fixed for 48 h in 10% neutral buffered formalin.

### Immunohistochemistry and microscopy of cryosections and paraffin-embedded sections

For immunohistochemistry (IHC) analysis, frozen rat tissues were initially fixed with 4% paraformaldehyde (PFA) in PBS. Fixed samples from human nerve tissue and rat peripheral nerves, DRG and spinal cord tissues were either i) embedded in optimal cutting temperature (OCT) cryo-compound (Tissue-Tek, Sakura), sectioned into 10 µm tissue cryosections, subsequently thaw mounted and stored at −20°C until further use; or ii) dehydrated in a graded series of ethanol (70%-100%), cleared in xylol, paraffin-embedded and sectioned into 5 µm tissue sections. After OCT washing or paraffin washing and rehydration, cryosections were blocked with 10% donkey serum (Sigma-Aldrich) and paraffin-embedded sections were incubated with antigen retrieval buffers (10 mM sodium citrate, 0.05% Tween 20, pH 6; or 1 mM EDTA, pH 8), treated with H_2_O_2_ to quench endogenous peroxidase (in case of HRP-conjugated secondary antibodies) and blocked with 2.5% horse serum (Vector Laboratories) and 1x casein solution (Vector Laboratories). Primary antibodies and stains were then incubated at 4°C overnight, dissolved in blocking solution as specified in **Supplementary Table 1**: rabbit anti-KIF4A (MyBioSource, MBS2518989); mouse anti-neurofilament, pan-axonal (Covance, SMI-312R, RRID:AB_2314906); rabbit anti-GAP43 (Abcam, ab75810, RRID:AB_1310252); rat anti-MBP (Bio-Rad, aa82-87); mouse anti-S100 (Abcam, ab7852, RRID:AB_306138); mouse anti-GFAP (Sigma-Aldrich, MAB3402, RRID:AB_94844); mouse anti-neurofilament 200 (Sigma-Aldrich, N0142, RRID:AB_477257); mouse anti-neurofilament 160/200 (Sigma-Aldrich, N2912, RRID:AB_477262); mouse anti-CD34 (Santa Cruz Biotechnology, sc-74499, RRID:AB_1120394); mouse anti-NG2 (Santa Cruz Biotechnology, sc-53389, RRID:AB_784821); isolectin B4, FITC conjugated (Sigma-Aldrich, L2895). Tissue sections were subsequently washed with PBS and incubated with fluorophore-conjugated secondary antibodies in blocking solution [anti-rabbit IgG Alexa Fluor™ 594 (Thermo Fisher Scientific, A21207, RRID:AB_141637 and A11012, RRID:AB_2534079), anti-mouse IgG Alexa Fluor™ 488 (Thermo Fisher Scientific, A21202, RRID:AB_141607 and A11001, RRID:AB_2534069) and anti-rat IgG Alexa Fluor™ 488 (Thermo Fisher Scientific, A21208, RRID:AB_2535794); anti-rabbit IgG, F(abʹ)2 fragment-Cy3 and anti-mouse IgG, FITC (Sigma-Aldrich, C2306, RRID:AB_258792 and F0257, RRID:AB_259378)] or ready-to-use HRP-conjugated secondary antibodies [anti-mouse IgG and anti-rabbit IgG (Vector Laboratories, MP-7402, RRID:AB_2336528 and MP-7401, RRID:AB_2336529)] and washed again with PBS. Nuclei were counterstained with DAPI solution (1:10, Roche). After washing, fluorescence-labelled sections were immersed in 70% EtOH for 5 min, transferred to 0.3% Sudan Black (Sigma-Aldrich), and mounted with Immu-Mount^TM^ (Thermo Fisher). Fluorescence-labelled sections were photographed using either the fluorescence microscopes Axioplan 2 with AxioVision Rel 4.8 software, or an LSM 880 Meta confocal microscope (both from Carl Zeiss). Images were analyzed and edited with ImageJ 2.0.0. (NIH). HRP-labelled sections were revealed (approx. 1 min) using the Diaminobenzidine (DAB) Substrate Kit (Vector Laboratories, SK-4100). The enzymatic reaction was stopped with ddH_2_O and samples were counterstained with Harry’s hematoxylin (15 s) and washed in tap water. After staining, images were obtained with a Nikon eclipse microscope.

### Preparation and culture of primary rat neonatal Schwann cells

For proliferation assays, rat neonatal Schwann cells were collected and prepared according to Brockes *et al*. 1979^93^. Neonatal sciatic nerves were collected from Wistar rats, and the nerves were stripped of connective tissue and blood vessels before being cut into small pieces. These were placed in warm Dulbecco’s modified Eagle medium (DMEM) and centrifuged for 5 min at 720 x *g* to discard the supernatant. The pellet was digested with 10 mL of trypsin/collagenase for 60 min at 37 °C. The reaction was stopped by adding DMEM, followed by centrifugation at 405 x *g* for 5 min. The pellet was resuspended in medium and filtered with a 60 mm gaze filter. The filtrate was centrifuged at 405 x *g* for 5 min, resuspended in DMEM and incubated at 37 °C and 10% CO_2_. After 24 h, 10 mM of Arabinosyl Cytosine (Ara-C) was added for 48h, to prevent excessive fibroblast proliferation. After another 48-72h, two complement lysis were conducted, by a 20 min incubation at 37 °C with baby rabbit complement (Cedarlane) dissolved in DMEM, and 0.4 mL mouse anti-CD90 (Thy1) T11D7e antibody (AbD Serotec, MCA04G) (prediluted 1:1 in DMEM). Following a subsequentially centrifugation (5 min, 585 x *g*), cells were triturated in 5 mL Schwann cells proliferation medium supplemented with 100 mg/mL Bovine Pituitary Extract (BPE) (Thermo Fisher) and 2 μM Forskolin (Sigma-Aldrich), and then transferred into a poly-D-lysine-coated culture flask and incubated at 37 °C and 10% CO_2_. When cells were confluent, about 4-5 days afterwards, another complement lysis was performed. Purified Schwann cells cultures were expanded in DMEM supplemented with 10% fetal bovine serum (FBS), 100 pg/ml crude glial growth factor protein preparation and 2 µM Forskolin, for 4-6 weeks. Schwann cells were expanded in DMEM with 10% FBS, and 2 µM Forskolin for about 10 days. The cells were finally passaged (passage 2) and were harvested either as a proliferative subconfluent cell culture (approx. 40-50% cell density) or as a dense confluent cell layer (∼100%) that was left confluent (and thus growth-arrested) for at least 24 hours.

Immunocytochemistry of these two populations was performed via routine techniques^94^, using rabbit anti-KIF4A (MyBioSource, MBS2518989), mouse anti-S100 (Santa Cruz Biotechnology, sc-58839). Fluorophore-conjugated secondary antibodies (anti-rabbit IgG Alexa Fluor™ 488 (Invitrogen, A11008), anti-mouse IgG Alexa Fluor™ 594 (Invitrogen, A11005) were used as above for IHC, and nuclei were counterstained with VECTASHIELD® Antifade Mounting Medium with DAPI (Vector Laboratories, H-1200) or Hoechst 33258 (PolySciences, #09460). Images were taken with an Axioplan 2 epifluorescence or a LSM 880 Meta confocal microscope (Carl Zeiss).

For the shRNA *Kif4a* silencing assays, primary rat Schwann cells were isolated from the sciatic nerve of 3-day post-natal pups as previously described^95^. Briefly, after dissection, nerves were stripped from epineurium and perineurium, cut into small pieces and incubated in 1% collagenase and 10 mg trypsin inhibitor in HBSS with Glutamax (Gibco) for 30 minutes at 37°C with occasional shaking. After pelleting, cells were washed twice in DMEM with Glutamax (Gibco) supplemented with 10% FBS and 1% penicillin/streptomycin, triturated using a 1 mL pipette and plated onto PDL-coated plates. Fibroblast proliferation was inhibited by limited exposure to 10 μM cytosine arabinoside. Around day 5, primary SC were re-fed with medium containing 2 μM Forskolin and 5 ng/mL neuregulin. On day 7, non-confluent cells were subjected to complement-mediated cytolysis using Thy1.1 antibody (Serotec). After at least 1-day recovery, cells were re-plated onto PDL-coated slides in a 24 well-plate (0.15x10^6^ cells per well).

### *Kif4* short hairpin plasmids and transfection in Schwann cells

The short hairpin RNA (shRNA) against *Kif4* (sh2-Kif4-GFP, referred in this study as ‘shKif4’), was kindly provided by Dr. Mariano Bisbal from the INIMEC-CONICET-UNC, Argentina (plasmid map in **Supplementary Figure 5**)^40^. pCAGIG empty vector, used as transfection control, was created by *Spe* l digestion and re-ligation of the shKif4 plasmid. Plasmids were amplified by midi preparation from overnight cultures *of E. coli* transformed clones using the NucleoBond Xtra Midi Kit (Macherey-Nagel), according with the manufacturer’s instructions.

Transfection of the shKif4 was performed using LipoFectamine300 (Invitrogen), according to the manufacturer’s protocol. Cells were incubated with the transfection complexes in suspension, plated, and fixed or collected 48 hours after transfection. *Kif4a* downregulation after shRNA transfection was validated by real-time PCR.

### Ki-67 proliferation assay

Immunocytochemistry of these two populations was performed via routine techniques^94^, using rabbit anti-KIF4A (MyBioSource, MBS2518989), mouse anti-S100 (Santa Cruz Biotechnology, sc-58839) or rabbit anti-Ki-67 (Invitrogen, MA5-14520, and Abcam, #15580). Fluorophore-conjugated secondary antibodies (anti-rabbit IgG Alexa Fluor™ 488 (Invitrogen, A11008), anti-mouse IgG Alexa Fluor™ 594 (Invitrogen, A11005) were used as above for IHC, and nuclei were counterstained with VECTASHIELD® Antifade Mounting Medium with DAPI (Vector Laboratories, H-1200) or Hoechst 33258 (PolySciences, #09460). Images were taken with a Zeiss Axioplan 2 epifluorescence using Leica DMI6000 FFW Motorized inverted epifluorescence microscope (Advanced Light microscopy, i3S).

Ki-67-positive transfected (GFP-positive) cells were scored for every condition, using Cell counter plug-in for ImageJ (Fiji Version 2.3.0/1.53f). Only S-100-positive cells were considered. Proliferation rate was calculated by dividing ki-67-positive transfected cells by the total transfected cells, per condition. pCAGIG-GFP vector was used as control.

### RNA isolation and quantitative real-time PCR analyses

Total RNA extraction and subsequent DNase I treatment were performed using a modified protocol. Total RNAs were extracted according to the protocol described by Untergasser Lab (Untergasser A. “RNAprep—TRIzol combined with Columns” Untergasser’s Lab. 2008) using TRIzol^TM^ Reagent (Invitrogen) and RNeasy Mini spin columns (Qiagen). RNA yield was determined using a spectrophotometer ND 1000 (PeqLab, Biotechnologie).

Reverse transcription (RT) of total RNA to single-stranded cDNA was performed using the High-Capacity cDNA RT Kit (Thermo Fisher). Quantification of *Kif4a* expression was performed relative to the endogenous housekeeping genes Ornithine Decarboxylase (*Odc*) or Glyceraldehyde-3-Phosphate Dehydrogenase (*Gapdh*), using Power SYBR Green Master Mix and the 7900HT Fast real-time PCR System (Applied Biosystems). The *Rattus norvegicus Kif4a* (NM_001400930.1) and *Odc* (NM_012615.3) reference sequences were applied to design specific primer pairs (Primer Express 3.0 Software, Applied Biosystems): TCCGGGAGGATCCTAAGGAA (*Kif4a* fwd) and GTCCGAGGCAACTAGCACAGT (*Kif4a* rev), TTG ACC AGA TGA ACG GAG TG (*Krox20* fwd) and CAG AGA TGG GAG CGA AGC TA (*Krox20* rev), GTT CTC GCA GAC CAC CAT CT (*Oct6* fwd) and GTC TCC TCC AGC CAC TTG TT (*Oct6* rev), GGT TCC AGA GGC CAA ACA TC (*Odc* fwd) and GTT GCC ACA TTG ACC GTG AC (*Odc* rev), GAACGGGAAGCTCACTGGC (Gapdh fwd) and GCATGTCAGATCCACAACG (Gapdh rev). Expression fold was calculated by comparison with control samples. All measurements were performed in duplicate and at least three independent samples (3 animals or cell pools, per condition and time point) were analyzed per experimental group. Relative fold mRNA expression was calculated using the comparative threshold cycle method (ΔΔCt). PCR products specificity was determined and verified using the dissociation curve analysis feature.

### KIF4 protein-protein interaction network and enriched processes

Using STRING DB^96^ v11.5, a protein-protein interaction (PPI) network of known interactors of KIF4 for *Mus musculus* was obtained, using data from experiments and databases, with a confidence > 0.7. Genes up- and down-regulated in the distal stump at 7 days post sciatic nerve transection were extracted from the supplementary information of Arthur-Farraj *et al*^60^. The known interactors of KIF4 extracted from STRING DB were searched in this list of de-regulated genes obtained from Arthur-Farraj *et al*^60^, which allowed us to compile a list/network of genes whose expression is de-regulated in the peripheral nerve at 7 days post-injury^60^ and whose protein products are known to interact with KIF4 (STRING DB). Finally, the ClueGo and CluePedia v2.5.8 plugins from Cytoscape v3.9.0^97^ were used to search for enriched Gene Ontology (GO) terms for molecular functions, biological processes, and enriched Reactome Pathways, with experimental evidence.

### Statistical analyses

Data are presented as mean ± SD. Statistical analysis was performed with the GraphPad Prism 9.3.0 software (GraphPad Software, Inc.). For mRNA expression experiments of rat injury models, the differences were determined using the unpaired multiple T-test (Welch correction, without correction for multiple comparisons), comparing each condition with the Sham control normalized to 1. For the proliferation assays, it was used the unpaired T-test (Welch correction). Results were considered statistically significant at p < 0.05.

## ETHICAL STATEMENTS

Nerve injury experiments were conducted in adult Wistar rats (200-240 g) in compliance with a) the German Animal Protection Law and were performed with the permission of the State Office for Nature, Environment and Consumer Protection North Rhine-Westphalia (LANUV NRW, Recklinghausen, Germany; and b) the ethical committee of the University of Granada, Spain, following the European Union and Spanish Government guidelines for animal experimentation (EU Directive No. 63/2010, RD 53/2013), approval number 29/03/2022/052, grant FIS PI20-0318.

Experiments involving animals at i3S were subjected to prior approval by the local Ethics Committee and were conducted in strict compliance with European and Portuguese guidelines (Project Licence DGAV 002803/2021).

Additionally, healthy sensitive human nerves were donated by patients who were subjected to reconstructive craniofacial surgeries, after obtaining their written informed consent.

## FUNDING

This work was funded by the Portuguese Foundation for Science and Technology (FCT), Centro 2020 and Portugal2020 and the European Union (FEDER program), via the project GoBack (PTDC/CVT-CVT/32261/2017) and the doctoral grants of PDC (SFRH/BD/139974/2018) and BMS (2020.06525.BD). JPF work was funded by the FCT (SFRH/BPD/113359/2015 - program-contract described in paragraphs 4, 5, and 6 of art. 23 of Law no. 1001 57/ 2016, of August 29, as amended by Law no. 57/2017 of July 2019; PTDC/MED-NEU/1677/2021).The authors also acknowledge the support of the Institute of Biomedicine iBiMED (UIDB/4501/2020 and UIDP/4501/2020) and its LiM Bioimaging Facility – a PPBI node (POCI-01-0145-FEDER-022122); the support of the Research Commission of the Medical Faculty of the Heinrich-Heine-University (HHU) Düsseldorf, and of the Biologisch-Medizinisches Forschungszentrum (BMFZ) of HHU. The study was also supported by the Spanish “Plan Nacional de Investigación Científica, Desarrollo e Innovación Tecnológica, Ministerio de Economía y Competitividad (Instituto de Salud Carlos III)”, co-financed by the European Union (FEDER program), (FIS PI17-0393 and FIS PI20-0318); by the “Plan Andaluz de Investigación, Desarrollo e Innovación (PAIDI 2020), Consejería de Transformación Económica, Industria, Conocimiento y Universidades, Junta de Andalucía, España” (P18-RT-5059); and by the “Programa Operativo FEDER Andalucía 2014-2020, Universidad de Granada, Junta de Andalucía, España, co-financed by the European Union (FEDER program), (A-CTS-498-UGR18).

## ACKNOWLEDGEMENTS

The authors would like to acknowledge Jessica Schira and Brigida Ziegler for the preparation and supply of primary rat Schwann cells at Dusseldorf, and to Dr. Mariano Bisbal^40^ for having supplied shRNAs targeting *Kif4a*.

## COMPETING INTERESTS STATEMENT

The authors have no competing interests to declare.

## AUTHOR CONTRIBUTIONS

Conceptualization: SIV, FB, VC and HWM; Methodology: SIV, FB, PDC, BMS, JCA, JPF and VC; Formal analysis and investigation: PDC, BMS, JCA, VE, VC and SIV; Main supervision of the work: FB and SIV; Writing - original draft preparation: PDC, BMS, JCA and SIV; Writing - review and editing: all authors; Funding acquisition: SIV, FB, JR, VC and HWM; Resources: SIV, FB, JR, VC and HWM.

## Notes

### Competing Interest Statement

The authors have declared no competing interest.

## REFERENCES

1. Perez K, Novoa AM, Santamarina-Rubio E, et al. Incidence trends of traumatic spinal cord injury and traumatic brain injury in Spain, 2000-2009. Accid Anal Prev 2012; 46: 37–44.

2. Apostolova I, Irintchev A, Schachner M. Tenascin-R Restricts Posttraumatic Remodeling of Motoneuron Innervation and Functional Recovery after Spinal Cord Injury in Adult Mice. J Neurosci 2006; 26: 7849–7859.

3. Liu H-N, Giasson BI, Mushynski WE, et al. AMPA receptor-mediated toxicity in oligodendrocyte progenitors involves free radical generation and activation of JNK, calpain and caspase 3. J Neurochem 2002; 82: 398–409.

4. Wang KC, Koprivica V, Kim JA, et al. Oligodendrocyte-myelin glycoprotein is a Nogo receptor ligand that inhibits neurite outgrowth. Nature 2002; 417: 941–944.

5. Oertle T, van der Haar ME, Bandtlow CE, et al. Nogo-A inhibits neurite outgrowth and cell spreading with three discrete regions. J Neurosci 2003; 23: 5393–406.

6. Schwab JM, Failli V, Chédotal A. Injury-related dynamic myelin/oligodendrocyte axon-outgrowth inhibition in the central nervous system. Lancet 2005; 365: 2055–2057.

7. McKeon RJ, Schreiber RC, Rudge JS, et al. Reduction of neurite outgrowth in a model of glial scarring following CNS injury is correlated with the expression of inhibitory molecules on reactive astrocytes. J Neurosci 1991; 11: 3398–411.

8. Vieira SI, Rebelo S, Esselmann H, et al. Retrieval of the Alzheimer’s amyloid precursor protein from the endosome to the TGN is S655 phosphorylation state-dependent and retromer-mediated. Mol Neurodegener 2010; 5: 40.

9. Barroca N, Marote A, Vieira SI, et al. Electrically polarized PLLA nanofibers as neural tissue engineering scaffolds with improved neuritogenesis. Colloids Surf B Biointerfaces 2018; 167: 93–103.

10. Carriel V, Garzón I, Campos A, et al. Differential expression of GAP-43 and neurofilament during peripheral nerve regeneration through bio-artificial conduits. J Tissue Eng Regen Med 2017; 11: 553–563.

11. Carriel V, Garzón I, Alaminos M, et al. Histological assessment in peripheral nerve tissue engineering. Neural Regen Res 2014; 9: 1657–1660.

12. Marote A, Barroca N, Vitorino R, et al. A proteomic analysis of the interactions between poly(L-lactic acid) nanofibers and SH-SY5Y neuronal-like cells. AIMS Mol Sci 2016; 3: 661–682.

13. Ahuja CS, Nori S, Tetreault L, et al. Traumatic Spinal Cord Injury—Repair and Regeneration. Neurosurgery 2017; 80: S9–S22.

14. Bosse F, Hasenpusch-Theil K, Küry P, et al. Gene expression profiling reveals that peripheral nerve regeneration is a consequence of both novel injury-dependent and reactivated developmental processes. J Neurochem 2006; 96: 1441–1457.

15. Carroll SL, Worley SH. Wallerian Degeneration☆. In: Reference Module in Neuroscience and Biobehavioral Psychology. Elsevier. Epub ahead of print 2017. DOI: 10.1016/B978-0-12-809324-5.02077-0.

16. Jessen KR, Mirsky R, Lloyd AC. Schwann Cells: Development and Role in Nerve Repair. Cold Spring Harb Perspect Biol 2015; 7: a020487.

17. Brück W. The role of macrophages in Wallerian degeneration. Brain Pathol 1997; 7: 741–52.

18. Shamash S, Reichert F, Rotshenker S. The cytokine network of Wallerian degeneration: tumor necrosis factor-alpha, interleukin-1alpha, and interleukin-1beta. J Neurosci 2002; 22: 3052– 60.

19. Stoll G, Müller HW. Nerve injury, axonal degeneration and neural regeneration: basic insights. Brain Pathol 1999; 9: 313–25.

20. Kaiser R, Haninec P. [Degeneration and regeneration of the peripheral nerve]. Ceskoslov Fysiol 2012; 61: 9–14.

21. Bosse F. Extrinsic cellular and molecular mediators of peripheral axonal regeneration. Cell Tissue Res 2012; 349: 5–14.

22. Jessen KR, Mirsky R. The Success and Failure of the Schwann Cell Response to Nerve Injury. Front Cell Neurosci; 13. Epub ahead of print 11 February 2019. DOI: 10.3389/fncel.2019.00033.

23. David S, Aguayo AJ. Axonal elongation into peripheral nervous system ‘bridges’ after central nervous system injury in adult rats. Science 1981; 214: 931–3.

24. Mason MRJ, Lieberman AR, Anderson PN. Corticospinal neurons up-regulate a range of growth-associated genes following intracortical, but not spinal, axotomy. Eur J Neurosci 2003; 18: 789–802.

25. Mason MRJ, Lieberman AR, Grenningloh G, et al. Transcriptional upregulation of SCG10 and CAP-23 is correlated with regeneration of the axons of peripheral and central neurons in vivo. Mol Cell Neurosci 2002; 20: 595–615.

26. Chaisuksunt V, Zhang Y, Anderson P., et al. Axonal regeneration from CNS neurons in the cerebellum and brainstem of adult rats: correlation with the patterns of expression and distribution of messenger RNAs for L1, CHL1, c-jun and growth-associated protein-43. Neuroscience 2000; 100: 87–108.

27. Yang P, Qin Y, Bian C, et al. Intrathecal delivery of IL-6 reactivates the intrinsic growth capacity of pyramidal cells in the sensorimotor cortex after spinal cord injury. PLoS One 2015; 10: e0127772.

28. Sun J, Ji Y, Liang Q, et al. Expression of Protein Acetylation Regulators During Peripheral Nerve Development, Injury, and Regeneration. Front Mol Neurosci 2022; 15: 888523.

29. Lee-Liu D, Moreno M, Almonacid LI, et al. Genome-wide expression profile of the response to spinal cord injury in Xenopus laevis reveals extensive differences between regenerative and non-regenerative stages. Neural Dev 2014; 9: 12.

30. Hui SP, Sengupta D, Lee SGP, et al. Genome wide expression profiling during spinal cord regeneration identifies comprehensive cellular responses in zebrafish. PLoS One 2014; 9: e84212.

31. Petrova V, Eva R. The Virtuous Cycle of Axon Growth: Axonal Transport of Growth-Promoting Machinery as an Intrinsic Determinant of Axon Regeneration. Dev Neurobiol 2018; 78: 898– 925.

32. Ducommun Priest M, Navarro MF, Bremer J, et al. Dynein promotes sustained axonal growth and Schwann cell remodeling early during peripheral nerve regeneration. PLoS Genet 2019; 15: e1007982.

33. Gumy LF, Chew DJ, Tortosa E, et al. The kinesin-2 family member KIF3C regulates microtubule dynamics and is required for axon growth and regeneration. J Neurosci 2013; 33: 11329–45.

34. Belrose JL, Prasad A, Sammons MA, et al. Comparative gene expression profiling between optic nerve and spinal cord injury in Xenopus laevis reveals a core set of genes inherent in successful regeneration of vertebrate central nervous system axons. BMC Genomics 2020; 21: 540.

35. Powers J, Rose DJ, Saunders A, et al. Loss of KLP-19 polar ejection force causes misorientation and missegregation of holocentric chromosomes. J Cell Biol 2004; 166: 991–1001.

36. Aizawa H, Sekine Y, Takemura R, et al. Kinesin family in murine central nervous system. J Cell Biol 1992; 119: 1287–1296.

37. Takemura R, Nakata T, Okada Y, et al. mRNA expression of KIF1A, KIF1B, KIF2, KIF3A, KIF3B, KIF4, KIF5, and cytoplasmic dynein during axonal regeneration. J Neurosci 1996; 16: 31–5.

38. Heintz TG, Heller JP, Zhao R, et al. Kinesin KIF4A transports integrin β1 in developing axons of cortical neurons. Mol Cell Neurosci 2014; 63: 60–71.

39. Peretti D, Peris L, Rosso S, et al. Evidence for the involvement of KIF4 in the anterograde transport of L1-containing vesicles. J Cell Biol 2000; 149: 141–52.

40. Bisbal M, Wojnacki J, Peretti D, et al. KIF4 mediates anterograde translocation and positioning of ribosomal constituents to axons. J Biol Chem 2009; 284: 9489–97.

41. Doherty P, Williams E, Walsh FS. A soluble chimeric form of the L1 glycoprotein stimulates neurite outgrowth. Neuron 1995; 14: 57–66.

42. Woolhead CL, Zhang Y, Lieberman AR, et al. Differential effects of autologous peripheral nerve grafts to the corpus striatum of adult rats on the regeneration of axons of striatal and nigral neurons and on the expression of GAP-43 and the cell adhesion molecules N-CAM and L1. J Comp Neurol 1998; 391: 259–73.

43. Zeng H, Yu B, Liu N, et al. Transcriptomic analysis of α-synuclein knockdown after T3 spinal cord injury in rats. BMC Genomics 2019; 20: 851.

44. Samejima K, Samejima I, Vagnarelli P, et al. Mitotic chromosomes are compacted laterally by KIF4 and condensin and axially by topoisomerase IIα. J Cell Biol 2012; 199: 755–770.

45. Zhu C, Jiang W. Cell cycle-dependent translocation of PRC1 on the spindle by Kif4 is essential for midzone formation and cytokinesis. Proc Natl Acad Sci U S A 2005; 102: 343–8.

46. Midorikawa R, Takei Y, Hirokawa N. KIF4 motor regulates activity-dependent neuronal survival by suppressing PARP-1 enzymatic activity. Cell 2006; 125: 371–83.

47. Willemsen MH, Ba W, Wissink-Lindhout WM, et al. Involvement of the kinesin family members KIF4A and KIF5C in intellectual disability and synaptic function. J Med Genet 2014; 51: 487–94.

48. Kokkonen H, Siren A, Määttä T, et al. Identification of microduplications at Xp21.2 and Xq13.1 in neurodevelopmental disorders. Mol Genet genomic Med 2021; 9: e1703.

49. Kalantari S, Carlston C, Alsaleh N, et al. Expanding the KIF4A-associated phenotype. Am J Med Genet A 2021; 185: 3728–3739.

50. Cho SY, Kim S, Kim G, et al. Integrative analysis of KIF4A, 9, 18A, and 23 and their clinical significance in low-grade glioma and glioblastoma. Sci Rep 2019; 9: 4599.

51. Wang X-B, Ma W, Luo T, et al. A novel primary culture method for high-purity satellite glial cells derived from rat dorsal root ganglion. Neural Regen Res 2019; 14: 339–345.

52. Lin G, Finger E, Gutierrez-Ramos JC. Expression of CD34 in endothelial cells, hematopoietic progenitors and nervous cells in fetal and adult mouse tissues. Eur J Immunol 1995; 25: 1508– 16.

53. Rezajooi K, Pavlides M, Winterbottom J, et al. NG2 proteoglycan expression in the peripheral nervous system: upregulation following injury and comparison with CNS lesions. Mol Cell Neurosci 2004; 25: 572–84.

54. Richard L, Topilko P, Magy L, et al. Endoneurial Fibroblast-Like Cells. J Neuropathol Exp Neurol 2012; 71: 938–947.

55. Dun X-P, Parkinson DB. Transection and Crush Models of Nerve Injury to Measure Repair and Remyelination in Peripheral Nerve. pp. 251–262.

56. Liu Z, Jin Y-Q, Chen L, et al. Specific Marker Expression and Cell State of Schwann Cells during Culture In Vitro. PLoS One 2015; 10: e0123278.

57. Nagarajan R, Le N, Mahoney H, et al. Deciphering peripheral nerve myelination by using Schwann cell expression profiling. Proc Natl Acad Sci U S A 2002; 99: 8998–9003.

58. Barrette B, Calvo E, Vallières N, et al. Transcriptional profiling of the injured sciatic nerve of mice carrying the Wld(S) mutant gene: identification of genes involved in neuroprotection, neuroinflammation, and nerve regeneration. Brain Behav Immun 2010; 24: 1254–67.

59. Arthur-Farraj PJ, Latouche M, Wilton DK, et al. c-Jun reprograms Schwann cells of injured nerves to generate a repair cell essential for regeneration. Neuron 2012; 75: 633–47.

60. Arthur-Farraj PJ, Morgan CC, Adamowicz M, et al. Changes in the Coding and Non-coding Transcriptome and DNA Methylome that Define the Schwann Cell Repair Phenotype after Nerve Injury. Cell Rep 2017; 20: 2719–2734.

61. de Curtis I. Intracellular Mechanisms for Neuritogenesis. Boston, MA: Springer US. Epub ahead of print 2007. DOI: 10.1007/978-0-387-68561-8.

62. Sheng L, Hao S-L, Yang W-X, et al. The multiple functions of kinesin-4 family motor protein KIF4 and its clinical potential. Gene 2018; 678: 90–99.

63. Salonen V, Röyttä M, Peltonen J. The effects of nerve transection on the endoneurial collagen fibril sheaths. Acta Neuropathol 1987; 74: 13–21.

64. Lewis GM, Kucenas S. Perineurial glia are essential for motor axon regrowth following nerve injury. J Neurosci 2014; 34: 12762–77.

65. Richard L, Védrenne N, Vallat J-M, et al. Characterization of Endoneurial Fibroblast-like Cells from Human and Rat Peripheral Nerves. J Histochem Cytochem 2014; 62: 424–435.

66. Geuna S. The sciatic nerve injury model in pre-clinical research. J Neurosci Methods 2015; 243: 39–46.

67. Schaeffer V, Meyer L, Patte-Mensah C, et al. Sciatic nerve injury induces apoptosis of dorsal root ganglion satellite glial cells and selectively modifies neurosteroidogenesis in sensory neurons. Glia 2010; 58: 169–80.

68. Archibald S, Shefner J, Krarup C, et al. Monkey median nerve repaired by nerve graft or collagen nerve guide tube. J Neurosci 1995; 15: 4109–4123.

69. Méchaly I, Bourane S, Piquemal D, et al. Gene profiling during development and after a peripheral nerve traumatism reveals genes specifically induced by injury in dorsal root ganglia. Mol Cell Neurosci 2006; 32: 217–29.

70. Ribeiro A, Balasubramanian S, Hughes D, et al. β1-Integrin cytoskeletal signaling regulates sensory neuron response to matrix dimensionality. Neuroscience 2013; 248: 67–78.

71. Fawcett JW. An integrin approach to axon regeneration. In: Eye (Basingstoke). 2017, pp. 206–208.

72. Previtali SC, Feltri ML, Archelos JJ, et al. Role of integrins in the peripheral nervous system. Prog Neurobiol 2001; 64: 35–49.

73. Wu J, Stoica BA, Faden AI. Cell Cycle Activation and Spinal Cord Injury. Neurotherapeutics 2011; 8: 221–228.

74. Aguayo AJ, David S, Bray GM. Influences of the glial environment on the elongation of axons after injury: transplantation studies in adult rodents. J Exp Biol 1981; 95: 231–40.

75. Castelnovo LF, Bonalume V, Melfi S, et al. Schwann cell development, maturation and regeneration: a focus on classic and emerging intracellular signaling pathways. Neural Regen Res 2017; 12: 1013–1023.

76. Glenn TD, Talbot WS. Signals regulating myelination in peripheral nerves and the Schwann cell response to injury. Curr Opin Neurobiol 2013; 23: 1041–1048.

77. Gomez-Sanchez JA, Pilch KS, van der Lans M, et al. After Nerve Injury, Lineage Tracing Shows That Myelin and Remak Schwann Cells Elongate Extensively and Branch to Form Repair Schwann Cells, Which Shorten Radically on Remyelination. J Neurosci 2017; 37: 9086–9099.

78. Mirsky R, Woodhoo A, Parkinson DB, et al. Novel signals controlling embryonic Schwann cell development, myelination and dedifferentiation. J Peripher Nerv Syst 2008; 13: 122–35.

79. Painter MW, Brosius Lutz A, Cheng Y-C, et al. Diminished Schwann cell repair responses underlie age-associated impaired axonal regeneration. Neuron 2014; 83: 331–343.

80. Martini R, Schachner M. Immunoelectron microscopic localization of neural cell adhesion molecules (L1, N-CAM, and myelin-associated glycoprotein) in regenerating adult mouse sciatic nerve. J Cell Biol 1988; 106: 1735–46.

81. Haney CA, Sahenk Z, Li C, et al. Heterophilic binding of L1 on unmyelinated sensory axons mediates Schwann cell adhesion and is required for axonal survival. J Cell Biol 1999; 146: 1173–84.

82. Loers G, Chen S, Grumet M, et al. Signal transduction pathways implicated in neural recognition molecule L1 triggered neuroprotection and neuritogenesis. J Neurochem 2005; 92: 1463–1476.

83. Moos M, Tacke R, Scherer H, et al. Neural adhesion molecule L1 as a member of the immunoglobulin superfamily with binding domains similar to fibronectin. Nature 1988; 334: 701–3.

84. Xu G, Nie D, Wang W, et al. Optic nerve regeneration in polyglycolic acid-chitosan conduits coated with recombinant L1-Fc. Neuroreport 2004; 15: 2167–72.

85. Samatov TR, Wicklein D, Tonevitsky AG. L1CAM: Cell adhesion and more. Prog Histochem Cytochem 2016; 51: 25–32.

86. Ide C. Peripheral nerve regeneration. Neurosci Res 1996; 25: 101–21.

87. Guseva D, Angelov DN, Irintchev A, et al. Ablation of adhesion molecule L1 in mice favours Schwann cell proliferation and functional recovery after peripheral nerve injury. Brain 2009; 132: 2180–2195.

88. Weiss SW, Nickoloff BJ. CD-34 is expressed by a distinctive cell population in peripheral nerve, nerve sheath tumors, and related lesions. Am J Surg Pathol 1993; 17: 1039–45.

89. Wu G, Zhou L, Khidr L, et al. A novel role of the chromokinesin Kif4A in DNA damage response. Cell Cycle 2008; 7: 2013–20.

90. Ribeiro B, Cruz B, de Sousa BM, et al. Cell therapies for spinal cord injury: a review of the clinical trials and cell-type therapeutic potential. Brain. Epub ahead of print 27 February 2023. DOI: 10.1093/brain/awad047.

91. Carriel V, Garrido-Gómez J, Hernández-Cortés P, et al. Combination of fibrin-agarose hydrogels and adipose-derived mesenchymal stem cells for peripheral nerve regeneration. J Neural Eng 2013; 10: 026022.

92. Chato-Astrain J, Campos F, Roda O, et al. In vivo Evaluation of Nanostructured Fibrin-Agarose Hydrogels With Mesenchymal Stem Cells for Peripheral Nerve Repair. Front Cell Neurosci 2018; 12: 501.

93. Brockes JP, Fields KL, Raff MC. Studies on cultured rat Schwann cells. I. Establishment of purified populations from cultures of peripheral nerve. Brain Res 1979; 165: 105–18.

94. de Sousa BM, Correia CR, Ferreira JAF, et al. Capacitive interdigitated system of high osteoinductive/conductive performance for personalized acting-sensing implants. NPJ Regen Med 2021; 6: 80.

95. Montani L, Bausch-Fluck D, Domingues AF, et al. Identification of new interacting partners for atypical Rho GTPases: a SILAC-based approach. Methods Mol Biol 2012; 827: 305–17.

96. Szklarczyk D, Morris JH, Cook H, et al. The STRING database in 2017: quality-controlled protein-protein association networks, made broadly accessible. Nucleic Acids Res 2017; 45: D362–D368.

97. Shannon P, Markiel A, Ozier O, et al. Cytoscape: a software environment for integrated models of biomolecular interaction networks. Genome Res 2003; 13: 2498–504.

