## Supplementary material for "Unexpected Kif4a functions in adult regeneration encompass a dual role in neurons and in proliferative repair Schwann cells"

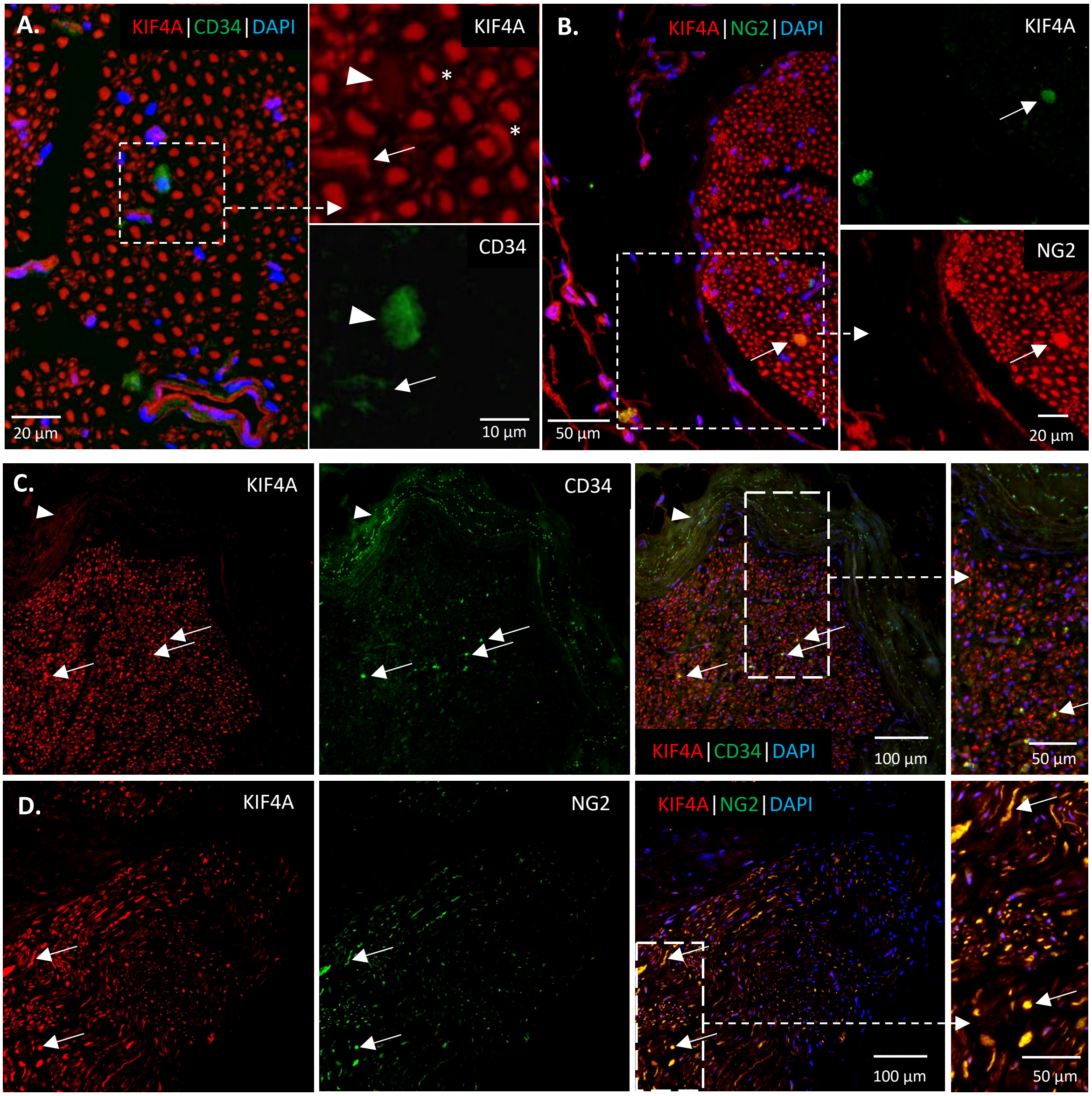


**Supplementary Figure 1. KIF4A distribution in CD34+ or NG2+ cells of rat and human uninjured peripheral nerves.** KIF4A (red) co-immunostaining with CD34 (green) (**A.** and **C.**) or NG2 (green) (**B.** and **D.**) in cross-sections of both rat (**A.** and **B.**) and human (**C.** and **D.**) sciatic nerves. KIF4A in axon-ensheathing Schwann cells (asterisk (*) in **A.**). Structures co-labelled with KIF4A and CD34 in rat and human sciatic nerves (arrows in **A.** and **C.**). Structures co-labelled with KIF4A and NG2 in rat and human sciatic nerves (arrows in **B.** and **D.**). Structures CD34-positive and KIF4A-negative, in both rat and human sciatic nerves (arrowheads in **A.** and **C.**).

**A. Nerve transection (with suture) at 7 dpi**

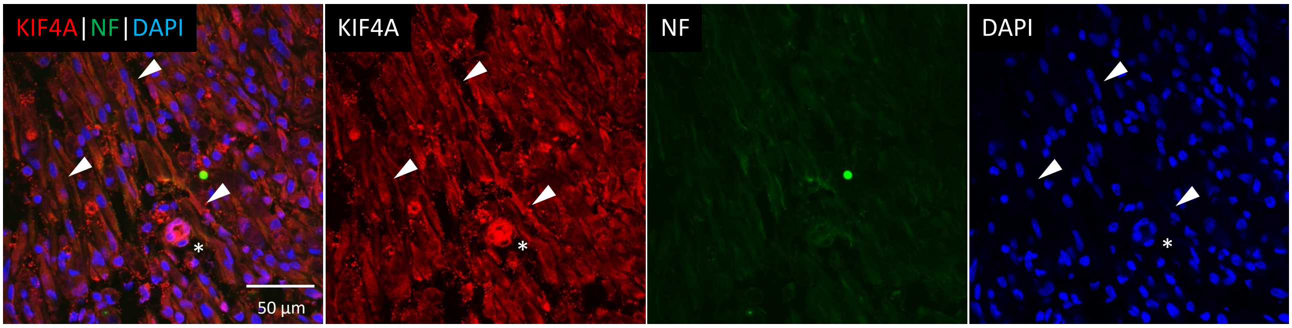


**B. Nerve transection (with suture) at 21 dpi**

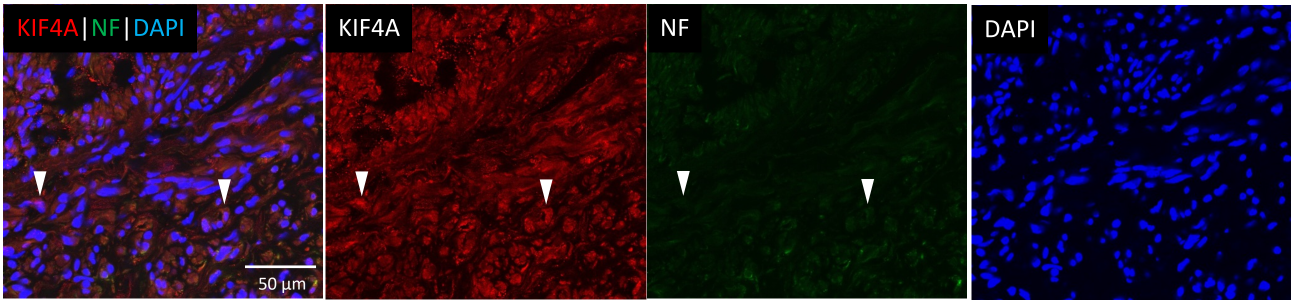


**C. Oct 6 mRNA levels with time post-injury**

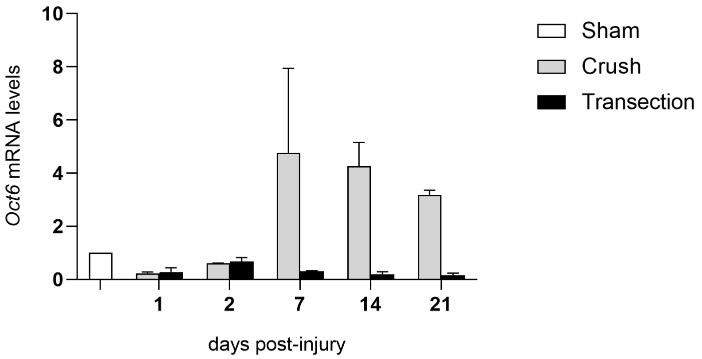


**Supplementary Figure 2. KIF4A distribution in sections of the distal stump of a transected and sutured sciatic nerve, and *Oct6* expression after injury.** KIF4A (in red) co-immunostaining with neurofilaments (NF, in green) in: **A.** a longitudinal section of a nerve collected 7 days post-injury (dpi); **B.** a cross-section of a nerve collected at 21 dpi. Note that NF staining in the distal stump disappears since axons were almost all lost due to Wallerian Degeneration. KIF4A immunostains various cellular structures (arrowheads) and what appears to be blood vessels (asterisks). **C.** Post-injury time-dependent expression profile of the Oct 6, marker of promyelinating Schwann cells (SCs), assessed by qRT-PCR. In the crush lesion condition, immature repair Schwann cells become promyelinating between 2 and 7 dpi, with this marker further slowly decreasing as SCs become myelinating ones. Contrary, in the transected with suture condition, the promyelinating phase is not achieved by the dedifferentiated SC. Values are presented as fold increases over mRNA levels of control sham-operated animals (taken as ‘1’). Results are presented as mean ± SD, n=2.

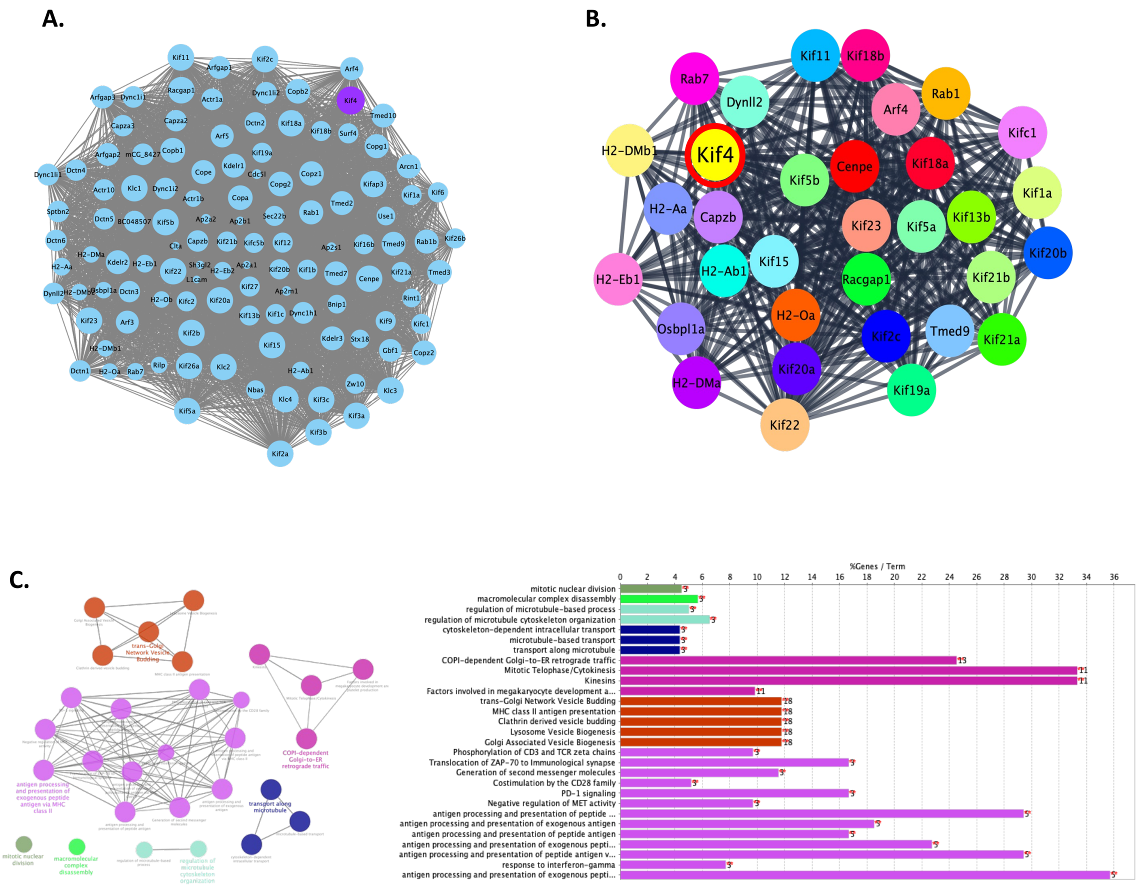


**D.**

| Function | Groups | Group Genes |
|---|---|---|
| mitotic nuclear division | Group 0 | Cenpe\|Kif11\|Kif18a |
| macromolecular complex disassembly | Group 1 | Capzb\|Kif19a\|Kif5b |
| regulation of microtubule cytoskeleton organization | Group 2 | Capzb\|Kif18a\|Kifc1 |
| transport along microtubule | Group 3 | Kif5a\|Kif5b\|Kifc1 |
| COPI-dependent Golgi-to-ER retrograde traffic | Group 4 | Arf4\|Capzb\|Cenpe\|Kif11\|Kif15\|Kif18a\|Kif20a\|Kif22\|Kif23\|Kif2c\|**Kif4**\|Kif5a\|Rab1a\|Racgap1\|Tmed9 |
| trans-Golgi Network Vesicle Budding | Group 5 | Cenpe\|Dynll2\|H2-Aa\|H2-Ab1\|H2-DMb1\|H2-Eb1\|H2Oa\| Kif11 \|Kif15\|Kif18a\|Kif22\|Kif23\|Kif2c\|**Kif4**\|Kif5a\|Osbpl1a\|Rab7\|Racgap1 |
| antigen processing and presentation of exogenous peptide antigen via MHC class II | Group 6 | H2-Aa\|H2-Ab1\|H2-DMb1\|H2-Eb1\|H2-Oa |

**Supplementary Figure 3. Kif4 PPI network and enriched pathways. A.** PPI network of Kif4 interactors in *Mus musculus* (obtained from String DB using data from experiments and databases, with confidence > 0.7). **B.** *Kif4* known interactors regulated in distal stump of mice transected sciatic nerves, at 7 days post-injury (from Arthur-Farraj *et al*. 2017). Node colours were randomly attributed. **C.** Enriched Reactome Pathways of the genes presented in **B. D.** Table with enriched pathways in **C.**, and the genes by group.

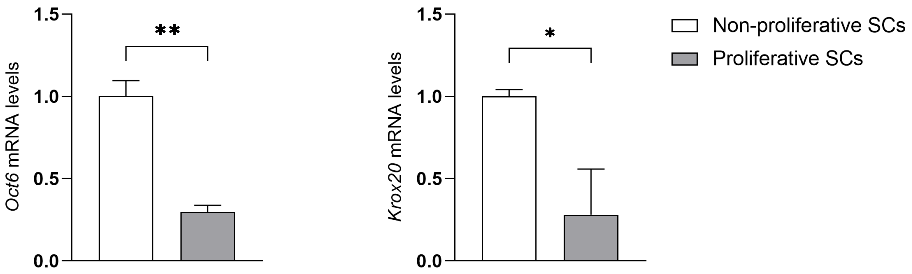


**Supplementary Figure 4. Characterization of the rat Schwann cells cultures used in terms of Oct6 and Krox20 expression.** Relative expression of *Oct6* and *Krox20* mRNAs, assessed by qRT-PCR, in the *in vitro* rat Schwann cells proliferation-state assay. This assay uses cells cultured at sub-confluent proliferative, or confluent density-arrested non-proliferative, conditions. Values are presented as fold increases over mRNA levels of control (taken as '1'). Results are presented as mean±SD, n=3. *, *p*-value <0.05; **, *p-value* <0.01, using the unpaired T-test with Welch correction.

**A.**

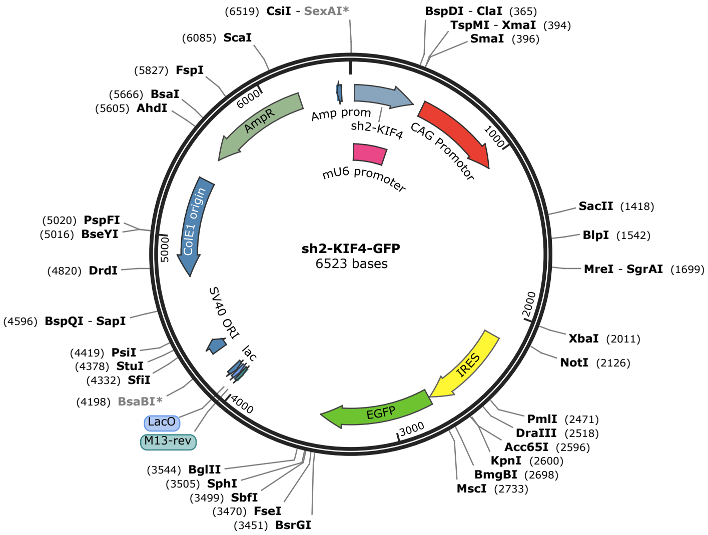


**B.**

| Plasmid | Antibiotic Resistance | Target sequence |
|---|---|---|
| Sh2-Kif4-GFP | Ampicilin | 5'-GGT GTT CCT GCA AAG GCT AAG CTT AGC CTT TGC AGG AAC ACC CTT TTT G-3' plus 5'-AAT TCA AAA AGG GTG TTC CTG CAA AGG CTA AGC TTA GCC TTT GCA GGA AAC C-3' |

**Supplementary Figure 5. Characterization of the shRNA used for Kif4a silencing. A.** Plasmid map of the anti-KIF4a shRNA used. **B.** Target sequence and plasmid information of the anti-Kif4a shRNA used.
This sh2-Kif4-GFP plasmid was provided by Dr. Mariano Bisbal (from Bisbal M, Wojnacki J, Peretti D, et al. KIF4 mediates anterograde translocation and positioning of ribosomal constituents to axons. *J Biol Chem* 2009; 284: 9489–97).

**Supplementary Table 1.** Technical information about the antibodies used for immunohistochemistry (IHC) and immunocytochemistry/immunofluorescence (IF) purposes.

| Antibody | Dilution/application | Supplier and Catalog number |
|---|---|---|
| **Rabbit anti-KIF4A** | 1:100 / IF and IHC | MyBioSource, MBS2518989 |
| **Mouse anti-neurofilament, pan-axonal** | 1:1000 / IF | Covance, SMI-312R |
| **Rabbit anti-GAP43** | 1:400 / IF and IHC | Abcam, ab75810 |
| **Rat anti-MBP** | 1:250 / IF | Bio-Rad, aa82-87 |
| **Mouse anti-S100** | 1:200 / IF | Abcam, ab7852 |
| **Mouse S-100 α/β chain (B32.1)** | 1:50/ IF | Santa Cruz Biotech., sc-58839 |
| **Mouse anti-GFAP** | 1:1000 / IF | Sigma-Aldrich, MAB3402 |
| **Mouse anti-Neurofilament 200** | 1:50 / IF | Sigma-Aldrich, N0142 |
| **Mouse anti-Neurofilament 160/200** | 1:500 / IF and IHC | Sigma-Aldrich, N2912 |
| **Mouse anti-CD34** | 1:50 / IF | Santa Cruz Biotech., sc-74499 |
| **Mouse anti-NG2** | 1:50 / IF | Santa Cruz Biotech., sc-53389 |
| **Rabbit anti-Ki67** | 1:250 / IF | Invitrogen, MA5-14520 |
| **Rabbit anti-Ki67** | 1:250 / IF | Abcam, #15580 |
| **Isolectin B4 (IB4 lectin)** | 1:20 / IF | Sigma-Aldrich, L2895 |
| **Donkey anti-rabbit IgG, Alexa Fluor™ 594** | 1:500 / IF | Thermo Fisher Scientific, A21207 |
| **Donkey anti-mouse IgG, Alexa Fluor™ 488** | 1:500 / IF | Thermo Fisher Scientific, A21202 |
| **Donkey anti-rat IgG, Alexa Fluor™ 488** | 1:500 / IF | Thermo Fisher Scientific, A21208 |
| **Goat anti-rabbit IgG, Alexa Fluor™ 594** | 1:300/ IF | Thermo Fisher Scientific, A11012 |
| **Goat anti-mouse IgG, Alexa Fluor™ 594** | 1:300/ IF | Thermo Fisher Scientific, A11005 |
| **Goat anti-rabbit IgG, Alexa Fluor™ 488** | 1:300/ IF | Thermo Fisher Scientific, A11008 |
| **Goat anti-mouse IgG, Alexa Fluor™ 488** | 1:300/ IF | Thermo Fisher Scientific, A11001 |
| **Sheep anti-rabbit IgG, F(ab')2 fragment-Cy3** | 1:100 / IF | Sigma-Aldrich, C2306 |
| **Goat anti-mouse IgG, FITC** | 1:100 / IF | Sigma-Aldrich, F0257 |
| **Horse anti-mouse IgG, HRP-conjugated** | pre-diluted / IHC | Vector Lab., MP-7402 |
| **Horse anti-rabbit IgG, HRP-conjugated** | pre-diluted / IHC | Vector Lab., MP-7401 |
